# Age-associated erosion of organ-specific endothelial programs compromises tissue function and resilience

**DOI:** 10.64898/2026.07.02.735775

**Authors:** Haruko Watanabe-Takano, Tomohiro Ishii, Tomohisa Hayakawa, Hitoshi Iuchi, Hitomi Matsuno, Eri Oguri-Nakamura, Kunihito Arai, Kei Yura, Michiaki Hamada, Daisuke Hishikawa, Shota Toyoshima, Mashito Sakai, Shimpei Higo, Masahiro Morishita, Hirotaka Ishii, Toru Tanaka, Sayo Horibe, Yoshiyuki Rikitake, Taichi Noda, Kimi Araki, Takashi Minami, Shinya Tanaka, Shigetomo Fukuhara

## Abstract

Endothelial cells (ECs) express organ-specific gene programs supporting tissue homeostasis and resilience. However, the mechanisms by which aging reshapes these organ-specific endothelial programs and how the resulting changes affect tissue homeostasis, resilience, and disease susceptibility remain largely unknown. Herein, we performed single-cell RNA sequencing of ECs harvested from five organs across the lifespan and found that aging progressively erodes organ-specific endothelial programs while inducing shared interferon-responsive and antigen-presentation programs across organs. Although vascular subtype identity and conserved capillary subset identity were mostly preserved, these organ-specific transcriptional programs were broadly attenuated with aging, indicating erosion of organ-specific endothelial identity to be a fundamental feature of endothelial aging. Importantly, these alterations were associated with declines in specialized EC functions, including alveolar barrier maintenance in the lung, scavenging activity in the liver, angiogenic capacity in the heart, and homeostatic programs in the kidneys and the brain, suggesting that age-related EC alterations compromise tissue homeostasis and resilience in multiple organs. Furthermore, we established a single-cell aging index for alveolar capillary ECs in mice and humans, revealing stress-associated endothelial activation to potentially be an intermediate state linking functional deterioration to cellular senescence, and also demonstrating marked heterogeneity in aging states even among ECs from animals of the same chronological age. Notably, alveolar capillary ECs exhibited progressive functional decline before reaching a senescent-like state, suggesting endothelial dysfunction to precede overt cellular senescence as the organism ages. Collectively, our findings establish progressive erosion of organ-specific endothelial programs as a central feature of vascular aging and provide a conceptual framework for elucidating how endothelial aging contributes to tissue dysfunction and reduced resilience across organs.

## Introduction

Endothelial cells (ECs) line the inner surfaces of blood vessels and form a vast vascular network that delivers oxygen and nutrients throughout the body, thereby sustaining life. Vascular aging has long been recognized as a key contributor to age-related disease development, and most studies published to date have focused on aging-associated endothelial dysfunction in the context of cardiovascular diseases, particularly atherosclerosis^1,2^. Furthermore, recent studies have shown that aging reduces capillary density in multiple organs, contributing to organ dysfunction and also increased mortality^3,4^. Collectively, these findings indicate that vascular aging research has been centered primarily on age-associated deterioration of vascular transport and perfusion functions.

However, blood vessels are now known to be more than passive conduits for blood transport. ECs actively regulate diverse tissue functions in an organ-specific manner and are increasingly recognized as being key regulators of tissue homeostasis and resilience^5–7^. Recent single-cell transcriptomic studies have revealed remarkable endothelial heterogeneity across organs^8,9^. Distinct EC populations establish specialized vascular architectural structures and express organ-specific transcriptional programs actively supporting tissue homeostasis and functions, including gas exchange and vascular barrier integrity in the lung, blood–brain barrier maintenance in the brain, scavenging and immune tolerance in the liver, filtration and tubular homeostasis in the kidneys, and metabolic support for cardiomyocytes.

These findings are consistent with recent single-cell transcriptomic analyses of individual organs that have identified core aging-associated endothelial signatures, including upregulation of inflammatory and interferon-responsive programs and downregulation of angiogenic programs^10–15^. However, despite these advances, how aging affects organ-specific endothelial programs across organs remains unclear and we do not as yet fully understand whether disruption of these programs contributes to impaired tissue function, reduced resilience, and increased susceptibility to age-associated diseases. A comprehensive multi-organ analysis of age-related endothelial alterations is therefore needed to determine how aging reshapes endothelial programs across organs and influences tissue homeostasis and resilience.

In this study, we performed single-cell RNA sequencing (scRNA-seq) of 258,574 ECs from five organs across the lifespans of mice and found aging to progressively erode organ-specific endothelial programs, thereby potentially compromising tissue homeostasis and resilience. In parallel, aging induced shared interferon-responsive and antigen-presentation programs, suggesting convergence toward common inflammatory states across organs. Furthermore, we established a single-cell aging index for alveolar capillary ECs in mice and humans, revealing stress-associated endothelial activation to potentially be an intermediate state in endothelial aging. Our observations also revealed substantial heterogeneity even among ECs from animals of the same chronological age. Notably, alveolar capillary ECs showed progressive functional decline before reaching a fully senescent state, suggesting loss of endothelial specialization to precede overt cellular senescence during aging.

## Results

### Comprehensive single-cell transcriptomic profiling of ECs across multiple organs in young and aged mice

To assess how aging reshapes organ-specific endothelial heterogeneity and function, we performed cross-organ scRNA-seq of ECs isolated from lung, kidney, liver, heart, and brain tissues of young (3 months) and aged (24–26 months) mice, including additional intermediate age groups for the lungs (6, 12, 15, and 17 months) and brain (12 months) (Fig. 1a). After quality control and batch integration using Harmony^16^, 258,574 high-quality ECs were retained for downstream analyses. ECs clustered predominantly by organ of origin rather than age (Fig. 1b and Extended Data Fig. 1a). Furthermore, Gene Ontology (GO) enrichment analysis of organ-enriched genes in young mice revealed functional programs aligned with the specialized physiological roles of each organ (Fig. 1c). As an example, pulmonary ECs were enriched for gene sets associated with cell-cell junction organization, extracellular matrix (ECM) binding, and ion channel activity, whereas hepatic ECs exhibited signatures related to lipoprotein metabolism, endocytosis, and immune-related processes. Similar organ-specific functional programs were also identified in other organs. These results indicate organ identity to be the principal determinant of EC transcriptional heterogeneity. Interestingly, a subset of organ-enriched genes overlapped with genes associated with parenchymal cell functions and this observation was validated in ECs by RNAscope and immunohistochemistry, suggesting specialized microenvironmental adaptation of ECs within each organ (Fig. 1d and Extended Data Fig. 1b,c).

**Fig. 1:**
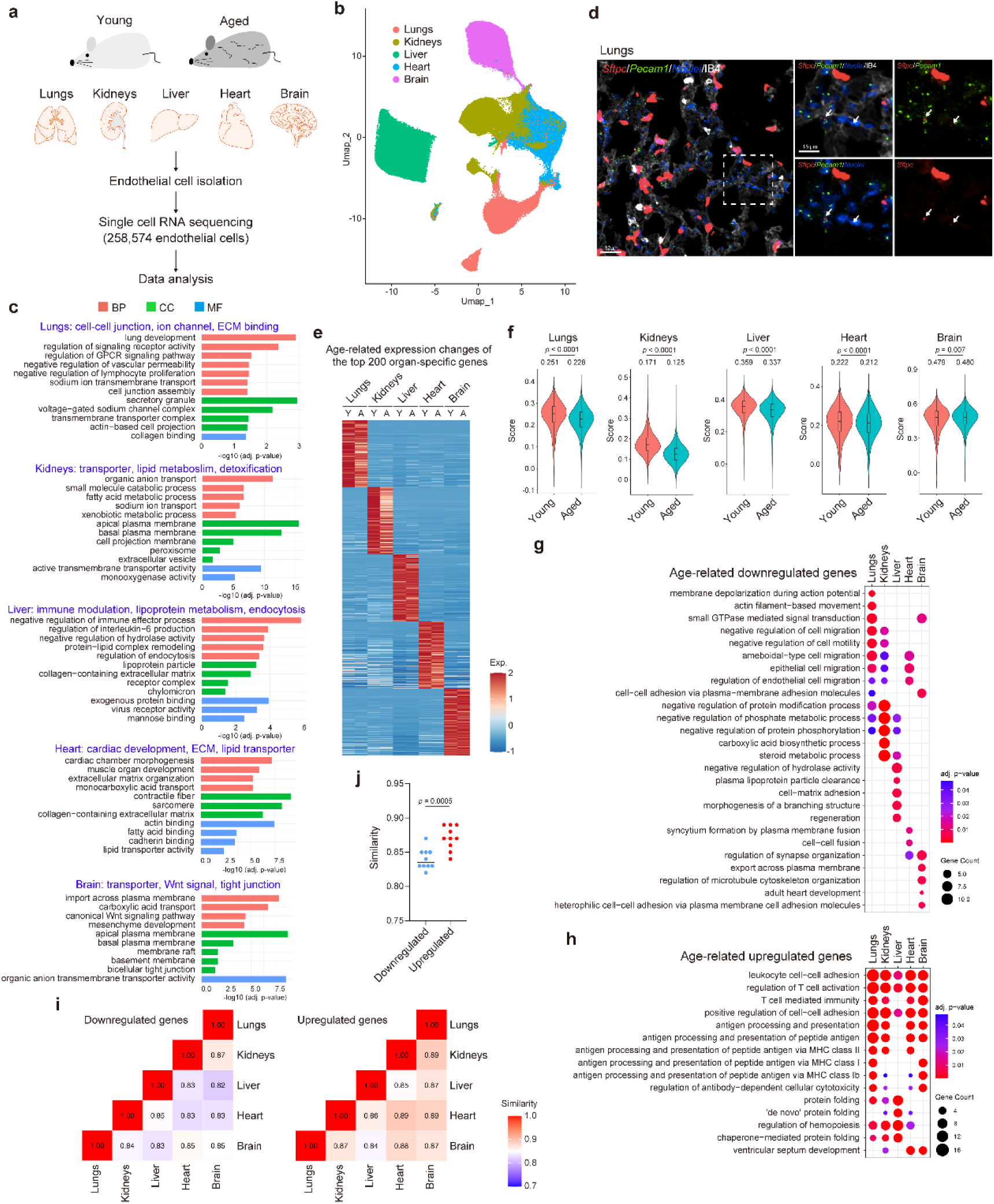
Comprehensive single-cell transcriptomic profiling of ECs across multiple organs in young and aged mice. **a**, Schematic of the experimental design. CD31⁺ ECs were isolated from five organs of young (3-month-old) and aged (24–26-month-old) mice and subjected to scRNA-seq, yielding 258,574 high-quality ECs. Additional intermediate age groups were included for the lungs (6, 12, 15, and 17 months) and brain (12 months). **b**, UMAP plot of ECs colored by tissue of origin. **c**, Bar plots showing GO terms enriched in the top 100 tissue-specific genes from each organ in young mice, categorized by biological process (BP, pink), cellular component (CC, green), and molecular function (MF, blue). x-axis, -log10(adjusted p-value). **d**, Representative confocal images of RNAscope in situ hybridization for *Sftpc* (red) and *Pecam1* (green) in alveoli from 12-week-old mice, counterstained with DAPI (nuclei, blue) and IB4 (white). Dashed box indicates the region enlarged in the right panels. Arrows indicate *Sftpc*-expressing *Pecam1*⁺ ECs. **e**, Heatmap showing normalized expression (z-score) of the top 200 organ-specific genes in ECs from each tissue of young (Y) and aged (A) mice. **f**, Violin plots overlaid with box plots showing module scores calculated from the top 200 tissue-specific genes identified in young mice, compared between young and aged mice. Boxes show median and interquartile range; median values are indicated above each plot. **g**,**h**, Dot plots showing GO (BP) terms enriched in the top 100 age-downregulated (**g**) and age-upregulated (**h**) genes across organs. Dot size, gene count; dot color, adjusted p-value. **i**, Heatmaps showing pairwise semantic GO (BP) similarity scores among organs for age-downregulated (left) and age-upregulated (right) genes, calculated using the GOSemSim package (Wang’s method, BMA strategy). **j**, Column scatter plots comparing inter-organ GO (BP) similarity scores for age-downregulated and age-upregulated genes as calculated in (**i**). Each dot represents a pairwise organ comparison. Scale bars, 30 μm (**d**) and 15 μm (enlarged panels in **d**). Wilcoxon rank-sum test (**f**). Unpaired two-tailed t test (**j**).

We next examined how aging changes these organ-specific programs and found many organ-enriched genes to be downregulated with aging in all organs except the brain (Fig. 1e,f). We also analyzed genes downregulated or upregulated with aging. GO terms associated with downregulated genes differed among organs and showed enrichment for organ-specific functional categories (Fig. 1g and Extended Data Fig.1d). In contrast, GO terms associated with upregulated genes were broadly shared, across organs examined, including functions such as antigen presentation and lymphocyte regulation (Fig. 1h and Extended Data Fig. 1e). Accordingly, inter-organ GO term similarity was greater for upregulated than for downregulated genes (Fig. 1i.j).

Collectively, these findings indicate that aging diminishes organ-specific endothelial identity while an immune-associated transcriptional program is commonly upregulated in all organs.

### Age-related attenuation of organ-specific gene expressions across vascular subtypes

We next assessed the impacts of aging on vascular subtypes. Clustering of ECs within each organ identified canonical vascular bed subtypes, including arterial, arteriolar, capillary, venular, and venous populations, as well as organ-specific specialized subtypes (Fig. 2a and Extended Data Fig. 2a,b). Each vascular subtype formed a continuum along the arterial–venous axis in the Uniform Manifold Approximation and Projection (UMAP) embedding, which reflected anatomical zonation from arteries to veins. Comparing young and aged mice revealed age-related shifts in capillary and venular populations in the lungs and capillary populations in the liver, whereas vascular subtype distributions were mostly preserved in other organs, suggesting that pulmonary and hepatic ECs are particularly susceptible to age-related alterations (Extended Data Fig. 2c).

**Fig. 2:**
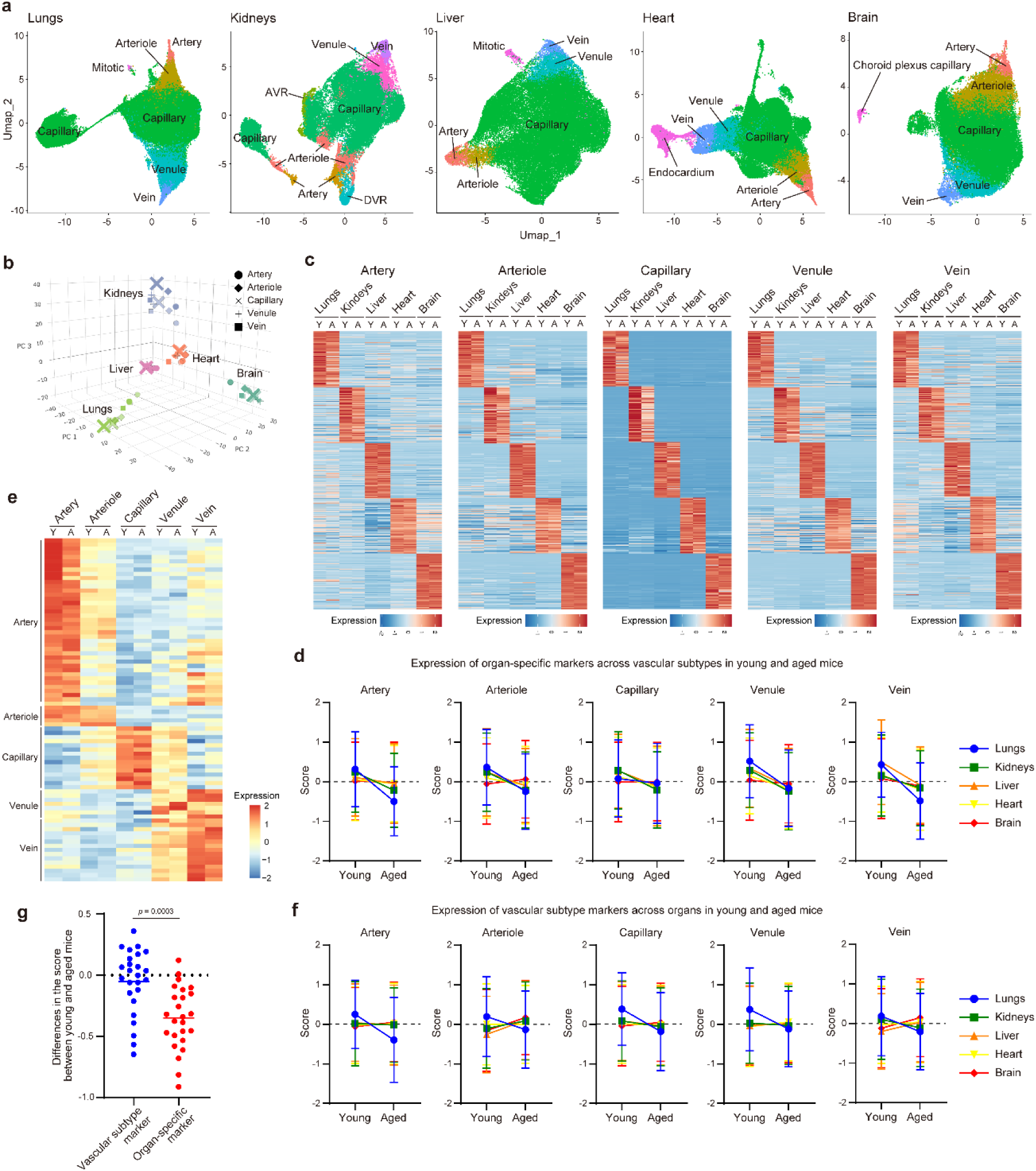
Age-related attenuation of organ-specific gene expression across vascular subtypes. **a**, UMAP plots of ECs from each organ, colored by vascular subtype. **b**, Three-dimensional PCA plot using the top 2,500 highly variable genes, showing clustering of vascular subtypes across five organs in young (dark) and aged (light) mice. **c**, Heatmaps showing normalized expression (z-score) of the top 100 tissue-specific genes across vascular subtypes from young (Y) and aged (A) mice. **d**, Z-scores for the expression levels of the top 300 organ-specific markers across vascular subtypes in young and aged mice. **e**, Heatmap showing normalized expression (z-score) of vascular subtype marker genes in young (Y) and aged (A) mice. **f**, Z-scores for the expression levels of vascular subtype markers across organs in young and aged mice. **g**, Differences in z-scores of vascular subtype markers (blue) and organ-specific markers (red) between young and aged mice. Unpaired two-tailed t test (**g**). Error bars, mean ± SD. (**d**, **f**)

Principal component analysis revealed ECs to primarily be segregated based on the organ of origin, rather than by vascular subtype identity (Fig. 2b). Among vascular subtypes, capillary ECs exhibited the highest degree of inter-organ heterogeneity, followed by arteriolar and venular subtypes, whereas arterial and venous populations exhibited relatively low heterogeneity. Consistently, each canonical vascular subtype displayed organ-specific transcriptional programs, with capillary populations having the highest degree of organ specificity (Fig. 2c). Organ-specific transcriptional programs within each vascular subtype were broadly attenuated with aging in all organs except the brain (Fig. 2c,d). Next, we examined the impacts of aging on vascular subtype identity. To this end, we identified marker genes defining vascular subtype identities across organs and found that their expressions within each vascular subtype were essentially preserved with aging (Fig. 2e). Expression scores of vascular subtype markers were maintained or even increased in most organs, while being predominantly decreased in the lungs (Fig. 2f). Consistently, the average expression scores of vascular subtype markers across all organs were mostly preserved as the mice aged, whereas organ-specific marker expressions were reduced (Fig. 2g). These results indicate that aging leads to selective deterioration of organ-specific transcriptional programs, while vascular subtype identities are largely preserved.

### Aging specifically disrupts organ-specific transcriptional programs in conserved capillary EC subsets

Capillary ECs are essential for maintaining organ function and showed the highest degree of organ specificity among vascular subtypes^8,9^. Thus, we next assessed the impacts of aging on capillary ECs in several organs. Clustering analysis identified multiple capillary EC subpopulations in each organ examined (Fig. 3a). Among them, Cap-Signal, Cap-AP-1, Cap-IFN, and Cap-Rpl,Rps, as well as Mitotic ECs, were conserved in all organs except the brain, where neither Cap-AP-1 nor Mitotic ECs were detected (Fig. 3a,b). In addition to Cap-IFN, a Cap-IFNg subset, characterized by type II IFN signaling, was observed in lungs and kidneys. We therefore analyzed the transcriptional similarity of these conserved EC populations across organs. Each conserved capillary EC subset exhibited similar gene expression patterns, varying minimally among the organs studied (Extended Fig. 3a). Consistently, subtype-specific similarity scores were higher within the same EC subset across organs than between distinct EC subsets (Fig, 3c,d). These findings indicate that conserved capillary EC subsets are shared across organs, as further validated by RNAscope and immunofluorescence analyses (Extended Fig. 3b-d).

**Fig. 3:**
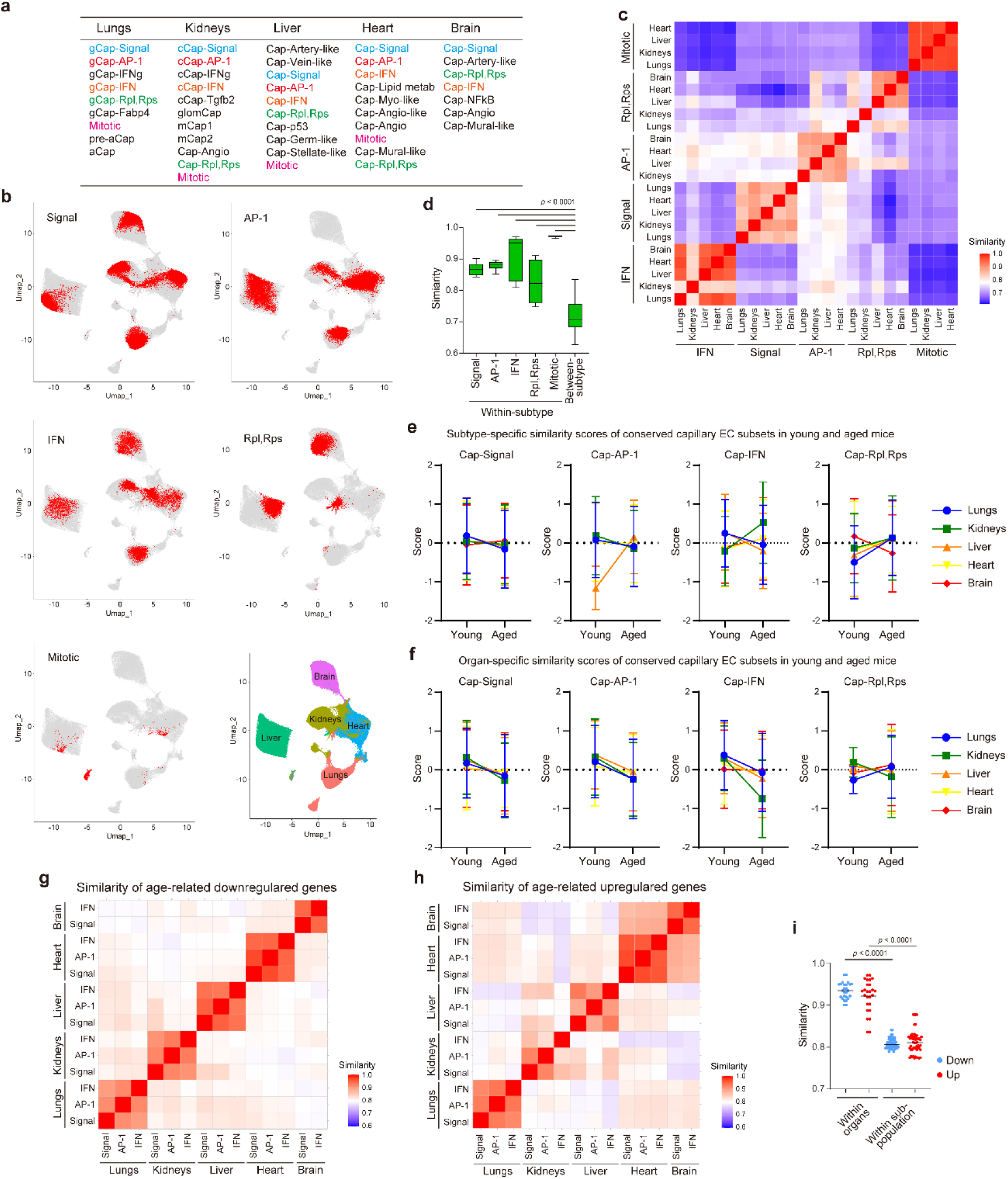
Aging preferentially disrupts organ-specific transcriptional programs in conserved capillary EC subsets. **a**, Capillary EC subpopulations identified in each organ. **b**, UMAP plots highlighting conserved capillary EC subsets (Signal, AP-1, IFN, Rpl,Rps, and Mitotic) across organs. **c**, Heatmap showing pairwise semantic GO (BP) similarity scores of subpopulation-specific marker genes across organs. **d**, Box plots showing GO (BP) similarity scores of subpopulation markers within the same subtype between organs (within-subtype) or within the same organ between subtypes. **e**, Z-scores for the expression levels of capillary subtype markers within conserved capillary EC subsets (Cap-Signal, Cap-AP-1, Cap-IFN, and Cap-Rpl,Rps) across organs in young and aged mice. **f**, Z-scores for the expression levels of organ-specific markers within conserved capillary EC subsets (Cap-Signal, Cap-AP-1, Cap-IFN, and Cap-Rpl,Rps) across organs in young and aged mice. **g**,**h,** Heatmap showing pairwise semantic GO (BP) similarity scores of the top 100 age-downregulated (**g**) and upregulated (**h**) DEGs across tissue–subpopulation combinations. **i**, Dot plots comparing GO (BP) similarity scores from (**g**) and (**h**), categorized into within-organ and within-subpopulation comparisons. Downregulated genes, blue; upregulated genes, red. Error bars, mean ± s.d. (**e**, **f**). One-way ANOVA with post hoc Dunnett’s test (**d**). One-way ANOVA with post hoc Tukey’s test (**i**).

We next analyzed aging-associated transcriptional alterations in conserved capillary EC subsets. Notably, subtype-specific transcriptional similarity was not consistently impacted by aging (Fig. 3e). In contrast, organ-specific transcriptional similarity was broadly attenuated with aging in Cap-Signal, Cap-AP-1, and Cap-IFN subsets in all organs examined except the brain, whereas the Cap-Rpl,Rps subset remained relatively preserved (Fig. 3f). We also analyzed aging-associated downregulated and upregulated genes in Cap-Signal, Cap-AP-1, and Cap-IFN subsets obtained from the organs studied. Importantly, aging-associated transcriptional alterations showed more similarity among different capillary EC subsets within the same organ than among the same EC subsets across organs (Fig. 3g-i). These results indicate that aging specifically compromises organ-specific transcriptional programs rather than subtype-specific capillary EC identity.

### Aging progressively erodes alveolar capillary EC homeostasis while promoting immune-activated states

In the lungs, capillary ECs form the blood–gas barrier together with alveolar epithelial cells, facilitating efficient gas exchange, and are composed of two distinct subtypes, aerocytes (aCap) and general capillary (gCap) ECs^17^. Clustering of lung capillary ECs identified two aCap populations and six gCap populations, together with a mitotic EC population (Fig. 4a,b). Among gCap ECs, the gCap-Signal and gCap-AP-1 subsets were the predominant populations in young mice, but were gradually replaced by the gCap-IFNg population with aging (Fig. 4c and Extended Fig. 4a). These three subsets shared age-downregulated genes associated with GO categories including angiogenesis, cell migration, actin filament organization, and cell–cell junction organization, suggesting ongoing loss of homeostatic functions with aging (Extended Fig. 4b,c). The aCap subset also decreased with aging and shared age-downregulated gene programs with the three gCap subsets. In contrast, all gCap and aCap subsets exhibited upregulation of gene programs associated with antigen processing and presentation, major histocompatibility complex class I (MHC-I) and class II (MHC-II) molecules, and leukocyte cell-cell adhesion (Extended Fig. 5a.b). These results raise the possibility that aging shifts alveolar capillary ECs from homeostatic states toward antigen-presenting states that might modulate T-cell responses.

**Fig. 4:**
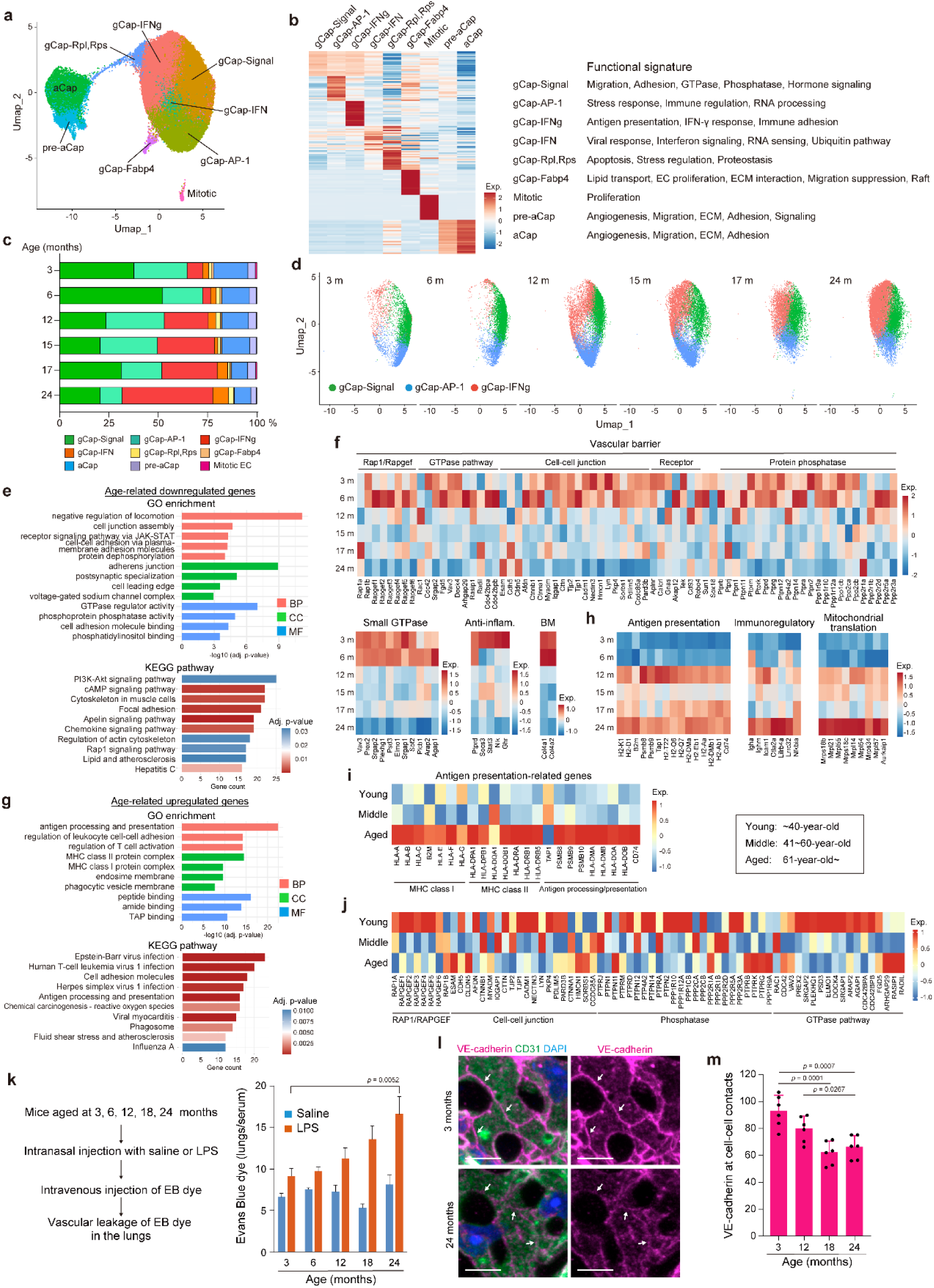
Aging progressively erodes alveolar capillary endothelial homeostasis while promoting immune-activated states. **a**, UMAP plot of alveolar capillary ECs with subpopulations indicated by colors. **b**, Heatmap showing the top 10 marker genes for each subpopulation identified in young mice, with functional signatures indicated on the right. Color scale, z-score. **c**, Proportional changes in alveolar capillary EC subpopulations across age groups. **d**, UMAP plots of tgCap ECs (gCap-Signal, gCap-AP-1, and gCap-IFNg) at each time point. **e**, GO terms (upper) and KEGG pathways (lower) enriched in the top 100 age-downregulated genes in tgCap ECs. (GO term) x-axis, -log10(adjusted p-value); BP, pink; CC, green; MF, blue. (KEGG pathway) x-axis, gene count; color, adjusted p-value. **f**, Heatmaps showing normalized expression (z-score) of vascular barrier-related genes (Rap1/Rapgef, GTPase pathway, cell-cell junction, receptor, and protein phosphatase), small GTPase, anti-inflammatory, and basement membrane (BM) genes in tgCap ECs at 3, 6, 12, 15, 17, and 24 months. **g**, GO terms (upper) and KEGG pathways (lower) enriched in the top 100 age-upregulated genes in tgCap ECs. x-axis, gene count; color, adjusted p-value. **h**, Heatmaps showing normalized expressions (based on z-scores) of antigen presentation-, immunoregulation-, and mitochondrial translation-related genes at the same time points as in (**f**). **i**,**j**, Heatmaps showing normalized expressions (based on z-scores) of antigen presentation-related genes (**i**) and functional genes (RAP1/RAPGEF, cell-cell junction, phosphatase, and GTPase pathway) (**j**) in alveolar capillary ECs from the Human Lung Cell Atlas (HLCA), stratified by age group (young, <40 years; middle-aged, 41–60 years; aged, >61 years) from healthy never-smoking donors. **k**, Experimental design for vascular permeability assessment (left). Evans Blue dye extravasation in the lungs, normalized to serum concentration, following intranasal LPS or saline administration in mice at 3, 6, 12, 18, and 24 months of age (right). **l**, Representative confocal images of alveolar capillary ECs stained for VE-cadherin (magenta), CD31 (green), and DAPI (blue) in 3- and 24-month-old mice. Arrows indicate VE-cadherin at cell-cell junctions. **m**, Quantification of VE-cadherin fluorescence intensity at cell-cell junctions across age groups. Scale bars, 10 μm (**l**). Error bars, mean ± s.d. (**k**). One-way ANOVA with Tukey’s post hoc test (**k**, **m**).

Since the gCap-Signal and gCap-AP-1 subsets showed progressive replacement by the gCap-IFNg population with aging, we collectively defined these populations as transitional gCap (tgCap) ECs and analyzed their age-associated state transitions (Fig. 4d). Alveolar capillary ECs establish a strong vascular barrier via tight endothelial junctions, thereby limiting pulmonary edema during inflammation and infection^18^. As we reported, the small GTPase Rap1 is essential for maintaining alveolar vascular barrier integrity^19,20^. tgCap ECs exhibited broad age-related downregulation of genes associated with vascular barrier integrity, including genes involved in Rap1 signaling, cell junction assembly, protein dephosphorylation, and cyclic AMP (cAMP) signaling, suggesting ongoing deterioration of alveolar vascular barrier function with aging (Fig. 4e,f and Extended Fig. 5c). Furthermore, genes involved in alveolar vascular function and homeostasis, including small GTPase signaling, basement membrane (BM) organization, anti-inflammatory responses, and tissue repair, showed uniform downregulation with aging (Fig. 4f and Extended Fig. 5d). Conversely, genes associated with antigen processing and presentation and leukocyte cell-cell adhesion, as well as pathways related to viral infection, immunoregulation, mitochondrial translation, the electron transport chain, and glutathione conjugation, were upregulated in tgCap ECs (Fig. 4g,h and Extended Fig. 5e,f). Taken together, these findings indicate aging to be associated with progressive erosion of endothelial programs involved in alveolar capillary homeostasis while immune-activated EC states in tgCap ECs are actually enhanced.

Next, we turned our attention to whether similar age-related alterations in alveolar capillary ECs occur in human subjects by reanalyzing publicly available human scRNA-seq data from the Human Lung Cell Atlas (HLCA), a Human Cell Atlas flagship resource^21^. Alveolar capillary ECs from aged individuals showed upregulation of antigen presentation-related genes as compared with those from young and middle-aged subjects (Fig. 4i). In contrast, many, though not all, genes associated with alveolar capillary function and homeostasis were downregulated in the aged group (Fig. 4j). Our findings suggest that age-associated alterations of alveolar capillary ECs, characterized by loss of homeostatic programs and immune activation, are conserved between mice and humans.

### Age-related decline in alveolar capillary barrier integrity increases susceptibility to acute lung injury

Next, we aimed to determine whether age-related downregulation of vascular barrier-related genes increases susceptibility to acute lung injury (ALI). To achieve this objective, we intranasally administered lipopolysaccharide (LPS) to mice at 3, 6, 12, 18, and 24 months of age and assessed pulmonary vascular permeability (Fig. 4k). Basal pulmonary permeability differed minimally among the age groups. However, LPS-induced pulmonary vascular leakage progressively worsened with aging, suggesting age-dependent impairment of vascular barrier resilience. Consistent with this observation, junctional localization of vascular endothelial-cadherin in alveolar capillary ECs also decreased progressively with aging (Fig. 4l,m and Extended Fig. 5g). We previously demonstrated that mice with endothelial-specific attenuation of Rap1 signaling have increased susceptibility to pulmonary vascular leakage despite showing no apparent differences from wild-type mice under basal conditions^20^. Collectively, these findings indicate that age-related deterioration of alveolar capillary barrier integrity contributes to greater susceptibility to ALI.

### A single-cell aging index reveals progressive functional deterioration in alveolar capillary ECs

Age-related transcriptional alterations in tgCap ECs did not strictly correlate with chronological age (Fig. 4f,h), suggesting heterogeneity in aging among individual ECs. Thus, we endeavored to quantify EC aging states at the single-cell level using aging-associated upregulated genes, including interferon-responsive and mitochondrial stress-associated genes (Fig. 5a). Based on their expression levels, we calculated an interferon (IFN) score for each tgCap EC. The IFN score was highly correlated with expressions of the genes used for score calculation as well as with other aging-associated upregulated genes (Fig. 5a and Extended Fig. 6a). Furthermore, the IFN score progressively increased along the age-associated transition trajectory of tgCap ECs when we conducted UMAP embedding (Fig. 5b,c and Extended Fig. 6b). Importantly, the expressions of genes involved in alveolar capillary function progressively decreased as IFN scores rose (Fig. 5d). Consistently, most genes associated with alveolar capillary function and homeostasis exhibited gradual downregulation at lower IFN scores followed by sharp declines at higher IFN scores (Fig. 5e and Extended Fig. 6c). These results indicate IFN scores to have the potential to serve as a quantitative aging index reflecting ongoing functional deterioration in individual tgCap ECs. Despite the average aging index increasing with chronological age, substantial intercellular heterogeneity persisted within each age group (Fig. 5f), indicating that EC aging is not a uniform process but proceeds heterogeneously even among cells from animals of the same chronological age.

**Fig. 5:**
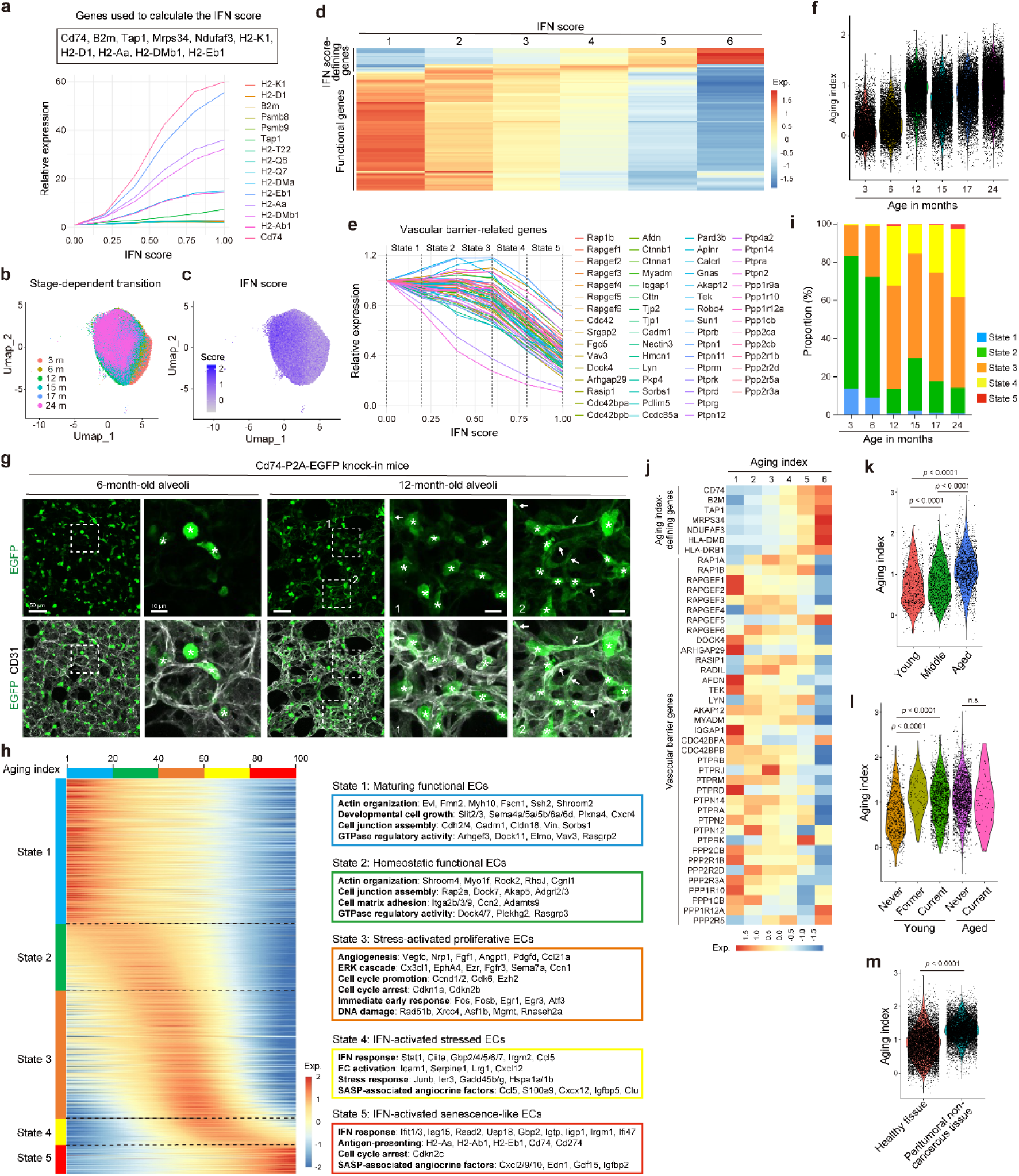
A single-cell aging index reveals progressive functional deterioration in alveolar capillary ECs. **a**, Relative expressions of IFN score-defining genes in tgCap ECs plotted against the IFN score. **b**,**c**, UMAP plots of tgCap ECs, with mouse age (**b**), indicated by color, or IFN score (**c**). **d**, Heatmap showing normalized expressions (based on z-scores) of IFN score-defining genes (upper) and functional genes (vascular barrier, anti-inflammatory, and basement membrane) (lower) in tgCap ECs, categorized into six groups based on IFN score. **e**, Relative expressions of vascular barrier-related genes in tgCap ECs plotted against the IFN score. **f**, Dot plots overlaid on violin plots showing the aging index of individual tgCap ECs across age groups. **g**, Representative confocal immunofluorescence images of alveoli from Cd74-P2A-EGFP knock-in mice at 6 months (left) and 12 months (right) of age, stained for GFP (green) and CD31 (white). Dashed boxes indicate enlarged regions. Asterisks and arrows indicate GFP-positive non-ECs and GFP-positive ECs, respectively. **h**, Heatmap of genes dynamically regulated along the aging index trajectory, identified by tradeSeq and ordered by peak expression (top bar, 0–100 bin). tgCap ECs are stratified into five states of 20 bins each: State 1, maturing functional ECs; State 2, homeostatic functional ECs; State 3, stress-activated proliferative ECs; State 4, IFN-activated stressed ECs; State 5, IFN-activated senescence-like ECs. Representative GO terms and associated genes for each state are shown. Color scale, z-score. **i**, Proportions of tgCap ECs in States 1–5 across age groups. **j**, Heatmap showing normalized expressions (based on z-scores) of aging index-defining genes (upper) and vascular barrier-related genes (lower) in human alveolar capillary ECs, categorized into six groups based on aging index. **k**, Dot plots overlaid on violin plots showing the aging index in human alveolar capillary ECs from young (<40 years), middle-aged (41–60 years), and aged (>61 years) healthy never-smoking donors. **l**, Dot plots overlaid on violin plots showing the aging index in human alveolar capillary ECs from young (<40 years) and aged (>61 years) donors stratified by smoking status (never, former, and current smokers). **m**, Dot plots overlaid on violin plots showing the aging index in human alveolar capillary ECs from healthy donors and peritumoral non-cancerous lung tissue of lung cancer patients. Scale bars, 50 μm (**g**) and 10 μm (enlarged panels in **g**). Kruskal–Wallis test followed by Bonferroni-corrected Dunn’s multiple comparison test (**k**-**m**).

To confirm the presence of functionally deteriorated alveolar capillary ECs in aged lungs, we generated *Cd74*-P2A-EGFP knock-in mice, as Cd74, the MHC-II invariant chain, was markedly induced along the EC aging trajectory, which enabled us to visualize these cells *in vivo* (Fig. 5a). In 6-month-old mice, strong EGFP signals were detected in macrophages and type II alveolar epithelial cells, whereas signals were minimal or even absent in alveolar capillary ECs (Fig. 5g). In contrast, 12-month-old mice exhibited EGFP signals in a subset of alveolar capillary ECs, frequently appearing as clusters rather than isolated cells (Fig. 5g). These findings suggest that alveolar capillary ECs, with functional deterioration, accumulate as clusters during the processes of aging.

### Stress-associated endothelial activation may drive age-related deterioration of alveolar capillary ECs

With the aim of characterizing the biological processes underlying age-related transition of tgCap ECs, we applied tradeSeq to identify genes dynamically regulated along the aging index trajectory (Fig. 5h). Based on the aging index, tgCap ECs were stratified into five sequential states representing progressive endothelial aging. Each of these states has a distinct transcriptional program. In States 1 and 2, genes associated with cell junction assembly, GTPase regulation, and actin organization, were expressed, observations consistent with functionally active ECs. State 1 cells were further enriched for the developmental growth-related genes, suggesting ECs undergoing maturation. In contrast, State 3 cells had upregulated genes associated with both EC activation (angiogenesis, ERK signaling, and cell cycle progression) and cellular stress responses (cell cycle arrest, immediate early response, and DNA damage), suggesting a stress-activated EC state. State 4 cells showed further upregulation of IFN-responsive genes and senescence-associated secretory phenotype (SASP)-associated angiocrine factors, whereas in State 5 cells genes associated with antigen presentation and cell cycle arrest were highly expressed, consistent with senescent-like EC states. These findings suggest a progressive transition from homeostatic to stress-activated, IFN-responsive, and ultimately senescent-like EC states, indicating stress-associated EC activation to possibly drive age-related deterioration of alveolar capillary ECs.

We next examined age-related changes in the proportions of State 1–5 cells and found that functional State 2 ECs predominated in young mice (3–6 months) (Fig. 5i). However, the proportion of State 2 ECs progressively diminished with aging, whereas stress-activated ECs in States 3 and 4 had gained dominance in 24-month-old mice.

Importantly, senescent-like State 5 ECs persisted as a minor population (∼ 3%) even in aged mice (Fig. 5i). Taken together with the findings that expressions of alveolar capillary functional genes fell from State 3 onward (Fig. 5e and Extended Fig. 6c) and that susceptibility to lung injury was already increased in 12-month-old mice (Fig. 4k), these results point to alveolar capillary ECs losing organ-specific functional resilience before reaching a fully senescent state.

### Environmental stress promotes premature aging of human alveolar capillary ECs

Using publicly available human scRNA-seq datasets^21^, we calculated IFN scores for human alveolar capillary ECs using genes related to antigen presentation and IFN signaling. Consistent with the data obtained from mice, higher IFN scores correlated with reduced expressions of alveolar capillary EC functional genes (Fig. 5j), supporting the usefulness of the IFN score as a conserved aging index. The average aging index increased with age, though there was substantial intercellular heterogeneity within each age group (Fig. 5k). In young individuals, alveolar capillary ECs from current and former smokers had higher aging indices than those from non-smokers, whereas aged individuals showed high aging indices regardless of smoking status (Fig. 5l). In addition, alveolar capillary ECs in peritumoral noncancerous lung tissue had higher aging indices than those in healthy lungs (Fig. 5m). These findings implicate environmental stress, in addition to chronological aging, in the promotion of premature aging of alveolar capillary ECs.

### Aging induces vascular bed-specific remodeling of renal capillary ECs

Renal function depends on highly specialized glomerular, cortical, and medullary capillary EC networks that regulate glomerular filtration, tubular transport, oxygen delivery, and osmotic balance^22^. We performed a clustering analysis which identified one glomerular capillary subset, five cortical capillary subsets, and two medullary capillary subsets together with three minor capillary subsets (Extended Fig. 7a.b).

In cortical capillary ECs, the cCap-Signal predominated in young mice and genes associated with small GTPase signaling, cell migration, and cell–cell junction organization were expressed, consistent with a homeostatic EC state (Extended Fig. 7b-d). We observed marked expansion of the immune-activated cCap-IFNg and fibrotic cCap-Tgfb2 subsets, whereas the cCap-IFN subset progressively diminished, with aging (Extended Fig. 7c,d). Across cortical capillary EC subsets, aging attenuated transcriptional programs associated with detoxification, metabolism, antioxidant responses, and transporter activities, while broadly inducing MHC-II-related genes and ER stress-response programs (Extended Fig. 7e-h). These findings suggest cortical capillary ECs from homeostatic states to shift toward dysfunctional, immune-activated, and stress-associated states with aging.

In glomerular capillary ECs, the glomCap subset exhibited age-related downregulation of endothelial structural and homeostatic genes together with upregulation of genes associated with TGF-β signaling, fibrosis, ECM remodeling, and ER stress responses, whereas MHC-related and IFN-responsive genes remained low during the processes of aging (Extended Fig. 7e-h). These findings are consistent with a transition of glomerular capillary ECs toward matrix-remodeling EC states.

In the medullary capillary subsets mCap1 and mCap2, aging attenuated ion transport-associated programs that are required for renal medullary homeostasis (Extended Fig. 7f). These subsets also downregulated renal homeostasis-associated programs while inducing the expression of MHC-II-related, lipid handling-associated, and stress-response genes. These findings suggest age-related remodeling of medullary capillary ECs to favor immune-activated and metabolically altered states.

### Aging remodels LSECs from homeostatic toward capillarized and pro-fibrotic states

Liver functions depend critically upon highly specialized hepatic sinusoidal ECs (LSECs), which regulate waste scavenging, lipid metabolism, immune tolerance, and molecular exchange between the blood and hepatocytes^23^. Clustering analysis identified nine capillary EC populations together with a mitotic subset (Fig. 6a,b). Gene expression analysis revealed that the Cap-Artery-like and Cap-Vein-like subsets corresponded, respectively to periportal zone 1 and pericentral zone 3 LSECs^24^. LSECs also included a homeostatic Cap-Signal subset and a Cap-Rpl,Rps subset enriched for ribosomal and oxidative phosphorylation-related genes. In addition, young livers contained a minor Cap-AP-1 population characterized by AP-1/immediate early-response, inflammatory, and stress-associated programs (Fig. 6b,c). However, this population gradually expanded with aging, whereas the Cap-Artery-like, Cap-Vein-like, and Cap-Signal subsets decreased (Fig.6c and Extended Fig. 8a). Consistently, c-Jun-positive LSECs were significantly increased in aged mice (Fig.6d and Extended Fig. 8b), supporting age-associated expansion of the Cap-AP-1 population.

**Fig. 6:**
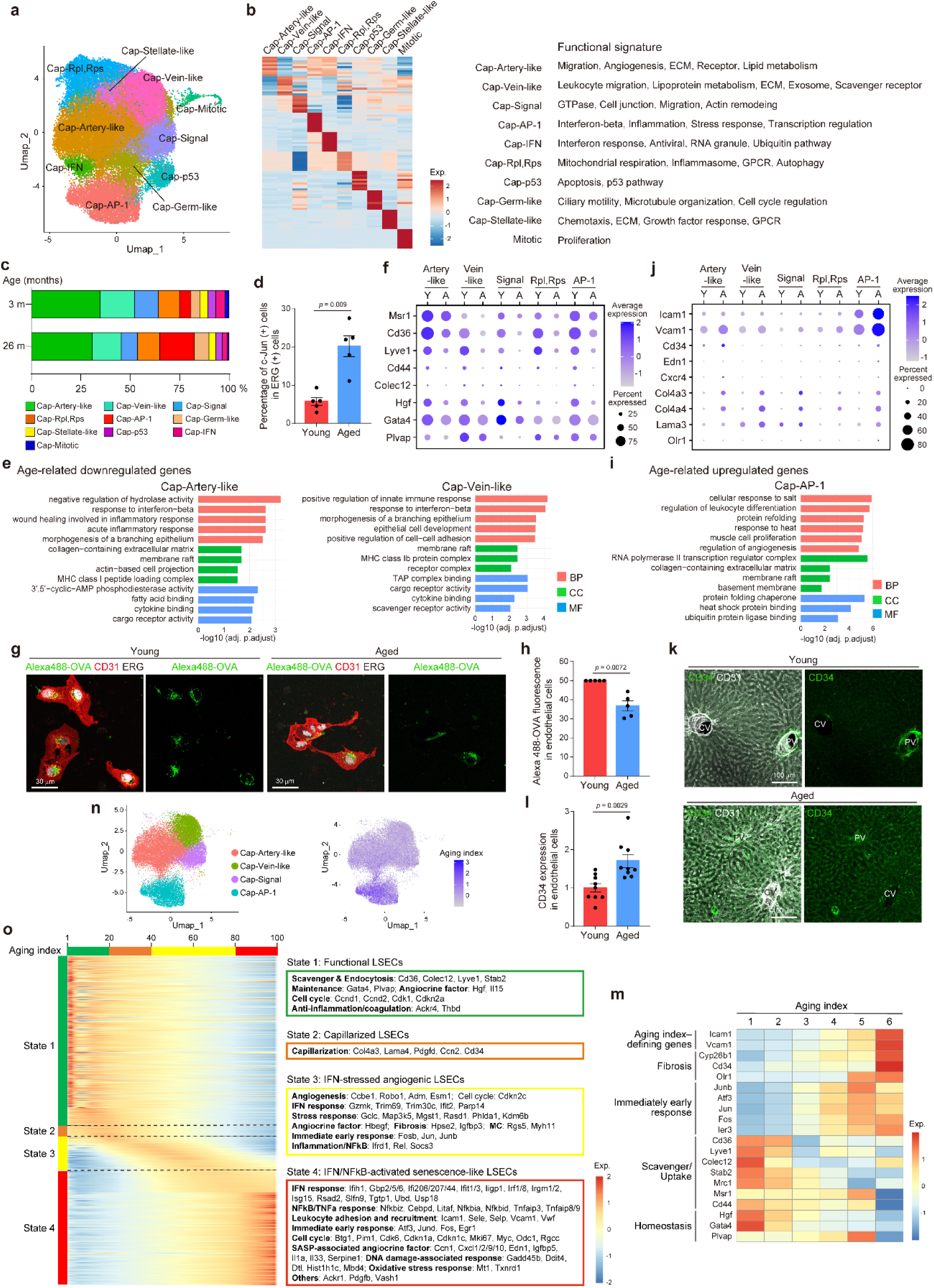
Aging remodels LSECs from homeostatic states toward capillarized and profibrotic states. **a**, UMAP plot of liver sinusoidal ECs (LSECs), with subpopulations indicated by coloer. **b**, Heatmap showing the top 10 marker genes for each subpopulation identified in young mice, with functional signatures indicated on the right. Color scale, z-score. **c**, Proportional changes in LSEC subpopulations in young (3-month-old) and aged (26- month-old) mice. **d**, Percentage of c-Jun-positive cells among ERG-positive ECs in the livers of young and aged mice. **e**, Bar plots showing GO terms enriched in the top 100 age-downregulated genes in Cap-Artery-like (left) and Cap-Vein-like (right) subpopulations. x-axis, -log10(adjusted p-value); BP, pink; CC, green; MF, blue. **f**, Dot plots showing expressions of scavenging-related genes (*Msr1*, *Cd36*, *Lyve1*, *Cd44*, and *Colec12*) and functional genes (*Gata4*, *Plvap*, and *Hgf*) in the indicated LSEC subpopulations of young and aged mice. Dot color, average expression; dot size, percentage of expressing cells. **g**, Representative confocal images of liver ECs isolated from young and aged mice following incubation with Alexa488-OVA (green). CD31 (red) and ERG (white) indicate ECs. **h**, Quantification of Alexa488-OVA fluorescence intensity within liver ECs from young and aged mice, as in (**g**). **i**, Bar plots showing GO terms enriched in the top 100 age-upregulated genes in the Cap-AP-1 subpopulation. x-axis, -log10(adjusted p-value); BP, pink; CC, blue; MF, green. **j**, Dot plots showing expressions of inflammatory genes (*Icam1*, *Vcam1*), capillarization-related genes (*Cd34*, *Col4a3*, *Col4a4*, *Lama3*), fibrosis-related genes (*Edn1*, *Olr1*), and angiogenesis-related genes (*Cxcr4*) in the indicated LSEC subpopulations of young and aged mice. Dot color, average expression; dot size, percentage of expressing cells. **k**, Representative confocal images of livers from young and aged mice stained for CD34 (green) and CD31 (white). CV, central vein; PV, portal vein. **l**, Quantification of CD34 fluorescence intensity within CD31-positive ECs from young and aged mice, as in (**k**). **m**, Heatmap showing normalized expressions (based on z-scores) of aging index-defining genes, immediate early response genes, fibrosis-related genes, scavenger/uptake-related genes, and homeostasis-related genes in major LSEC subpopulations (Cap-Artery-like, Cap-Vein-like, Cap-Signal, and Cap-AP-1), categorized into six groups based on aging index. **n**, UMAP plots of major LSEC subpopulations (Cap-Artery-like, Cap-Vein-like, Cap-Signal, and Cap-AP-1), with subpopulation (left), indicated by color, or aging index (right). **o**, Heatmap of genes dynamically regulated along the aging index trajectory, identified by tradeSeq and ordered by peak expression (top bar, 0–100). LSECs are stratified into four states: State 1 (1-20 bins), functional LSECs; State 2 (21-40 bins), capillarized LSECs; State 3 (41-80 bins), IFN-stressed angiogenic LSECs; State 4 (81-100 bins), IFN/NF-κB-activated senescence-like LSECs. Representative GO terms and associated genes for each state are shown. Color scale, z-score. Scale bars, 30 μm (**g**) and 100 μm (**k**). Error bars, mean ± s.e.m. (**d**, **h**, **l**). Unpaired two-tailed t test (**d**, **h**, **l**).

Our next analysis focused on aging-associated transcriptional alterations in LSECs. Genes involved in scavenging activity (*Msr1*, *Cd36*, *Lyve1*, *Cd44*, and *Colec12*) were broadly downregulated in all LSEC subsets with aging (Fig. 6e,f and Extended Fig. 8c). Consistently, uptake of Alexa488-labeled Ovalbumin (OVA) was reduced in aged LSECs as compared with young LSECs (Fig. 6g,h), indicating age-associated deterioration of LSEC scavenging functions. The expressions of *Gata4*, *Plvap*, and *Hgf*, which regulate LSEC specification, formation of fenestrae, and hepatic regeneration, respectively, were also reduced with aging (Fig. 6f)^25–27^. In parallel, MHC-I-associated antigen presentation genes were downregulated in multiple subsets (Fig. 6e and Extended Fig. 8c), suggesting tolerogenic immune programs in aged LSECs to be impaired. These findings point to an apparent progressive deterioration of specialized LSEC programs during aging.

Furthermore, aging induced expressions of genes associated with capillarization, angiogenesis, inflammation, and fibrosis in all LSEC subsets, though primarily in a small fraction of cells within each subset (Fig. 6i,j and Extended Fig. 8d). Consistently, expression of the capillarization marker CD34 was significantly increased in aged LSECs (Fig. 6k,l). Because capillarization disrupts specialized sinusoidal vascular architecture and promotes hepatic fibrosis^28^, these findings suggest that aging remodels LSECs toward capillarized and pro-fibrotic states. In parallel, AP-1/immediate early-response genes and chaperone-mediated protein folding genes were broadly upregulated in all LSEC subsets examined (Fig. 6i and Extended Fig. 8d), suggesting stress-associated and proteostatic remodeling as the organism ages. Collectively, these results indicate that aging remodels LSECs from specialized homeostatic states to favor capillarized, stress-associated, and pro-fibrotic EC states.

To investigate the cellular transitions underlying age-related deterioration of LSECs, we established an LSEC aging index applicable to the major populations (Cap-Artery-like, Cap-Vein-like, Cap-Signal, and Cap-AP-1) based on *Icam1* and *Vcam1* expression. The aging index showed positive correlations with fibrosis-associated genes (*Cyp26b1*, *Cd34*, and *Olr1*) and AP-1/immediate early-response genes, while being inversely correlating with genes involved in LSEC scavenging and homeostasis (Fig. 6m), indicating that the index captures functional deterioration of LSECs. Among the major populations examined, the Cap-AP-1 subset had the highest aging index (Fig. 6n), suggesting Cap-AP-1 LSECs to represent the most marked deterioration of the LSEC state during aging.

Based on the aging index, we stratified LSECs into four sequential states and analyzed genes enriched in each state (Fig. 6o). State 1 cells expressed LSEC functional and homeostatic genes, consistent with a homeostatic EC state. State 2 cells showed upregulation of capillarization-associated genes, whereas in State 3 cells further induction of the expressions of angiogenesis-, stress response-, and inflammation-associated genes was noted, again suggesting progressive transition toward activated and pro-fibrotic states during aging. Finally, genes associated with DNA damage, oxidative stress responses, and SASP-associated angiocrine factors were markedly upregulated in State 4 cells, observations consistent with senescence-like LSEC states. These results indicate capillarization- and angiogenesis-associated EC activation to possibly trigger stress-associated deterioration of LSECs as cells age

### Aging impairs angiogenic and regenerative programs in cardiac capillary ECs

Cardiac capillary networks support the high metabolic demands of the heart through continuous oxygen and nutrient delivery to cardiomyocytes^29^. Clustering analysis identified nine cardiac capillary EC subsets together with a mitotic EC subset (Fig. 7a,b). Two major subsets, Cap-Signal and Cap-AP-1, were detected in tissues from both young and aged mice (Fig. 7a-c and Extended Fig. 9a). The Cap-Signal subset expressed homeostatic genes associated with small GTPase signaling, cell–cell junction organization, and focal adhesion, consistent with maintenance of endothelial integrity and cardiac homeostasis. The Cap-AP-1 subset expressed AP-1/immediate early-response genes while retaining lower levels of homeostatic gene expression, which would be consistent with a stress-responsive yet functionally maintained EC state.

**Fig. 7:**
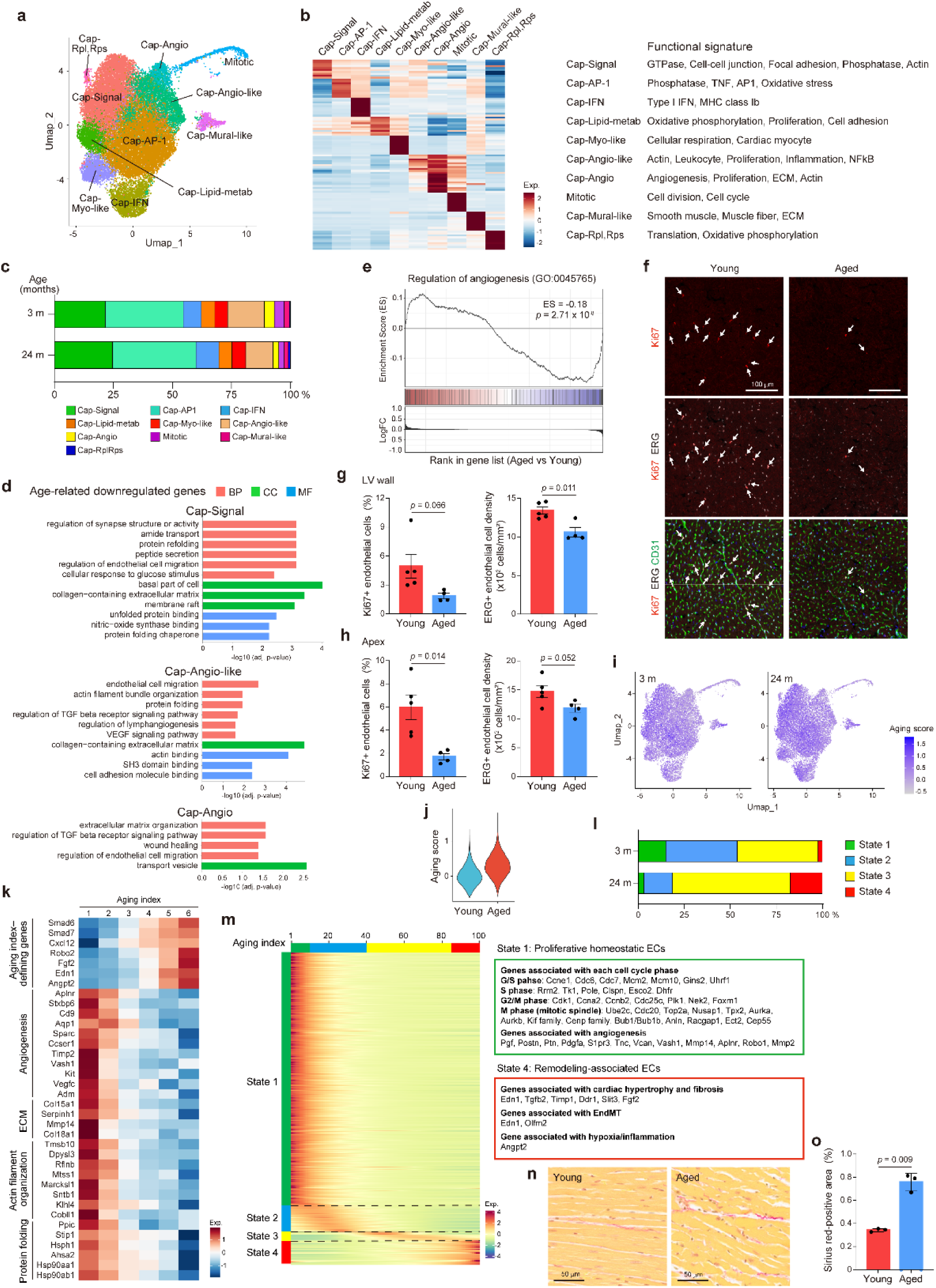
Aging impairs angiogenic and regenerative programs in cardiac capillary ECs. **a**, UMAP plot of cardiac capillary ECs, with subpopulations indicated by color. **b**, Heatmap showing the top 10 marker genes for each subpopulation identified in young mice, with functional signatures indicated on the right. Color scale, z-score. **c**, Proportional changes in cardiac capillary EC subpopulations in young (3-month-old) and aged (24-month-old) mice. **d**, Bar plots showing GO terms enriched in the top 100 age-downregulated genes in Cap-Signal, Cap-Angio-like, and Cap-Angio subpopulations. x-axis, -log10(adjusted p-value); BP, pink; CC, green; MF, blue. **e**, Gene set enrichment analysis (GSEA) plot for “regulation of angiogenesis” (GO:0045765) in aged as compared with young cardiac capillary ECs **f**, Representative confocal images of the left ventricular wall of hearts from young and aged mice stained for Ki67 (red), ERG (white), and CD31 (green). Arrows indicate Ki67- and ERG-double-positive ECs. **g**,**h**, Quantification of Ki67-positive ECs among ERG-positive ECs (left) and ERG-positive EC density (right) in the left ventricular wall (**g**) and apex (**h**) of young and aged mice. **i**, UMAP plots of cardiac capillary ECs, with aging index score indicated by color, in young (3-month-old) and aged (24-month-old) mice. **j**, Violin plots comparing aging index scores between young and aged cardiac capillary ECs. **k**, Heatmap showing normalized expressions (based on z-scores) of aging index-defining genes, angiogenesis-, ECM-, actin filament organization-, and protein folding-related genes in cardiac capillary ECs, categorized into six groups based on aging index. **l**, Proportions of cardiac capillary ECs in States 1–4 across age groups. **m**, Heatmap of genes dynamically regulated along the aging index trajectory, identified by tradeSeq and ordered by peak expression (top bar, 0–100). Cardiac capillary ECs are stratified into four states: State 1 (1–5 bins), proliferative homeostatic ECs; State 2 (6– 40 bins); State 3 (41–85 bins); State 4 (86–100 bins), remodeling-associated ECs. Representative GO terms and associated genes for each state are shown. Color scale, z-score. **n**, Representative light microscopy images of Sirius red staining in the heart tissues from young and aged mice. **o**, Quantification of Sirius red-positive areas in cardiac interstitial tissues from young and aged mice. Scale bars, 100 μm (**f**) and 50 μm (**n**). Error bars, mean ± s.d. (**g**, **h**, **o**). Welch’s t-test after the F-test indicated unequal variances (**g**, **h**, **o**).

In addition, the heart contained angiogenic and proliferative capillary populations, including the Cap-Angio-like, Cap-Angio, and mitotic ECs, reflected in a continuous trajectory on the UMAP embedding (Fig. 7a,b). Importantly, the proportion of these populations decreased from 23.9% to 16.2% with aging (Fig. 7c and Extended Fig. 9a). Furthermore, genes associated with angiogenesis, vascular endothelial growth factor signaling, and EC migration were broadly downregulated as aging progressed, across cardiac capillary EC subsets, and Gene set enrichment analysis further confirmed the suppression of angiogenic programs (Fig. 7d,e and Extended Fig. 9b). Consistently, aged mice exhibited reduced numbers of Ki67-positive ECs and total ECs in the left ventricular wall and apex as compared with young mice (Fig. 7f-h). Given that cardiac ECs have relatively high turnover rates and undergo angiogenic remodeling during physiological and pathological cardiac hypertrophy^30–32^, these findings suggest that aging impairs both the adaptive and the regenerative capacity of the cardiac vasculature.

In contrast, genes that were associated with antioxidant responses, haptoglobin signaling, and TGF-β signaling were upregulated with aging, in all cardiac capillary EC subsets (Extended Fig. 9c). MHC-I-related genes also showed upregulation with aging. These results provide evidence that aging may induce oxidative stress responses, fibrotic remodeling, and immune dysregulation in cardiac capillary ECs.

To investigate cardiac capillary EC aging, we established an aging index based on the expressions of 7 aging-associated genes: *Smad6*, *Smad7*, *Cxcl12*, *Fgf2*, *Robo2*, *Edn1*, and *Angpt2* (Fig. 7i-k). The aging index showed inverse correlation with angiogenesis-associated genes (Fig. 7k), indicating that the aging index captures the diminished angiogenic capacity of cardiac capillary ECs. In addition, genes involved in ECM organization, actin filament organization, and protein folding decreased as the aging index rose, suggesting deterioration of structural integrity and proteostasis during cardiac capillary EC aging.

Based on this novel aging index, we stratified cardiac capillary ECs into four sequential states and found the proportions of States 1 and 2 cells to fall as the organism aged, whereas those of States 3 and 4 increased (Fig. 7l,m). State 1 ECs expressed genes associated with cell cycle regulation and angiogenesis, while these programs showed an ongoing decrease with progression toward State 4 (Fig. 7m), indicating age-related impairment of regenerative and proliferative endothelial potential. In contrast, State 4 ECs showed high expressions of genes associated with cardiac hypertrophy, fibrosis, hypoxia, and inflammation (Fig. 7m). Aging has been reported to promote cardiac fibrosis and hypertrophy^33,34^, consistent with our observation of increased Sirius red staining of collagen fibers in cardiac tissues of aged mice compared with young mice (Fig. 7n,o). These findings suggest aging-altered cardiac capillary ECs to possibly contribute to pathological cardiac remodeling.

### Aging induces complex neurovascular remodeling in brain capillary ECs

Brain ECs form highly specialized blood–brain barrier (BBB) networks capable of tightly regulating molecular transport and neuronal homeostasis within the central nervous system^35,36^. Clustering analysis identified seven brain capillary EC subsets, with Cap-Signal, Cap-Artery-like, and Cap-Rpl,Rps constituting the major populations (Extended Fig. 10a,b). The Cap-Signal subset expressed genes related to small GTPase signaling, cell–cell junction organization, and insulin signaling, whereas the Cap-Artery-like subset expressed transporter- and ECM-associated genes, consistent with their roles in maintaining BBB integrity and function (Extended Fig. 10b). In contrast, the Cap-Rpl,Rps subset exhibited a metabolically active state with high expressions of genes associated with mitochondrial respiration and translation. Brain capillary EC subset composition remained relatively stable as aging progressed (Extended Fig. 10c,d), suggesting limited compositional remodeling.

Differentially expressed genes (DEG) analysis revealed aging to downregulate genes associated with BBB integrity and protection (*Gja1*^37^, *Cdh2*^38^, *Pcdh19*^39^, *Actn1*^40^, *Mt1*^41^), transport (*Abca1*^42^, *Slc38a5* ^43^), and anticoagulant function (*Thbd*^44^, *Serpine2*^45^) (Extended Fig. 10e,f), suggesting that BBB homeostatic programs deteriorate as the organism ages. *Htra1*, a causative gene for hereditary cerebral small vessel disease^46^, and *Flrt2*, a gene implicated in protection against vascular aging^47^, were also downregulated. Conversely, MHC-I-related genes and genes associated with inflammation, stress responses, bone morphogenic protein signaling, and vascular barrier dysfunction were upregulated as the organism aged (Extended Fig. 11a,b), suggesting that aging-induced remodeling of brain capillary ECs contributes to BBB dysfunction as well as neurovascular abnormalities.

Notably, as aging progressed, genes implicated in BBB dysfunction (*Sparc*^48^, *Pde4d*^49^, *Angptl4*^50^, *Adamts5*^51^) were downregulated simultaneously with the upregulation of genes involved in vascular barrier maintenance, BBB transport, and anti-inflammatory responses (Extended Fig. 10g and 11c), raising the possibility that aged brain ECs may mount a compensatory protective response against neurovascular stress. Collectively, these findings suggest that aging leads to complex neurovascular remodeling in brain capillary ECs, involving both maladaptive and compensatory responses.

### Aging enhances endothelial antigen presentation programs across organs

Aging was found to broadly induce interferon-associated pathways in several organs (Fig. 8a). We therefore investigated age-related changes in MHC-associated antigen presentation programs in ECs. Classical MHC-I-related genes are ubiquitously expressed in most cell types. Consistent with this observation, MHC-I-related genes were expressed in most ECs and further upregulated with aging in all organs examined except the liver (Fig. 8b). Accordingly, the proportion of MHC-I-positive ECs increased with aging (Fig. 8c), most prominently in the brain, where MHC-I-positive ECs rose from 7.6% in young to 25.2% in aged mice. Furthermore, H2-Q and H2-T families of non-classical MHC-Ib genes were upregulated with aging in capillary ECs of most organs (Fig. 8b). These findings point to aging as broadly enhancing endothelial MHC-I-associated antigen presentation programs across organs.

**Fig. 8:**
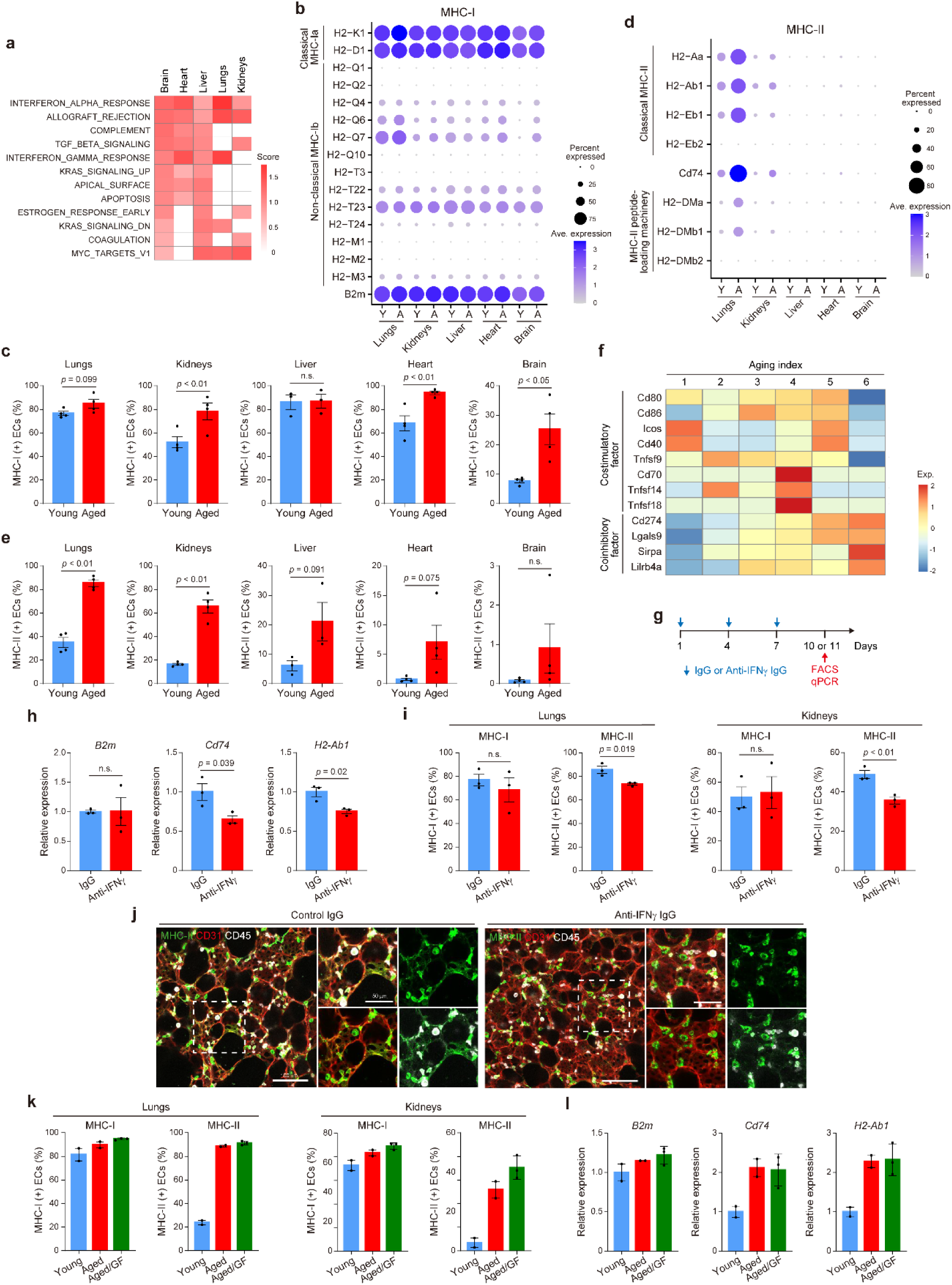
Aging enhances endothelial antigen presentation programs across organs. **a**, Heatmap showing age-related upregulated pathways identified by GSEA across organs. Color scale, normalized enrichment score (NES). **b**,**d**, Dot plots showing expression of MHC-I genes (**b**) and MHC-II-related genes (**d**) across organs in young (Y) and aged (A) mice. Dot color, average expression; dot size, percentage of expressing cells. **c**,**e**, Percentages of MHC-I-positive (**c**) and MHC-II-positive (**e**) ECs across organs from young and aged mice, as determined by flow cytometry. **f**, Heatmap showing normalized expressions (based on z-scores) of costimulatory and coinhibitory factor genes in tgCap ECs, categorized into six groups based on aging index. **g**, Experimental design for IFN-γ neutralization in aged mice. Control IgG or anti-IFN-γ IgG was administered every 3 days to 2-year-old mice, followed by flow cytometry, histological analysis, and qPCR. **h**, Relative mRNA expressions of *B2m*, *Cd74*, and *H2-Ab1* in lungs from aged mice treated with control IgG or anti-IFN-γ IgG, as determined by qPCR. **i**, Percentages of MHC-I-positive and MHC-II-positive ECs in lungs (left) and kidneys (right) of aged mice treated with control IgG or anti-IFN-γ IgG, as determined by flow cytometry. **j**, Representative confocal images of lungs from aged mice treated with control IgG or anti-IFN-γ IgG, stained for MHC-II (green), CD31 (red), and CD45 (white). Dashed boxes indicate enlarged regions. **k**, Percentages of MHC-I-positive and MHC-II-positive ECs in lungs (left) and kidneys (right) of young (12-week-old), aged (75-week-old), and germ-free aged (79-week-old) (Aged/GF) C57BL/6N mice, as determined by flow cytometry. **l**, Relative mRNA expressions of *B2m*, *Cd74*, and *H2-Ab1* in lungs of young, aged, and germ-free aged C57BL/6N mice, as determined by qPCR. Scale bars, 100 μm (**j**) and 50 μm (enlarged panels in **j**). Error bars, mean ± s.e.m. (**c**, **e**, **h**, **i**, **k**, **l**). Unpaired two-tailed t test (**c**, **e**, **h**, **i**): p values are indicated. One-way ANOVA with Tukey’s post hoc test (**k**, **l**).

MHC-II expression is generally restricted to highly specific antigen-presenting cells, such as dendritic cells, B cells, and macrophages. However, a considerable proportion of pulmonary and renal capillary ECs expressed MHC-II-related genes even in young mice, and their abundance and MHC-II gene expression both rose with aging (Fig. 8d). Consistently, the proportions of MHC-II-positive ECs significantly increased with aging in these organs (Fig. 8e). In the heart, liver, and brain, while MHC-II-expressing ECs were less abundant, flow cytometric analysis revealed a similar trend toward increased MHC-II-positive ECs as the mice aged. These findings suggest that aging promotes MHC-II-mediated antigen presentation by ECs across organs.

To explore the potential functional consequences of age-associated antigen presentation programs, we determined the expressions of costimulatory and coinhibitory molecules in tgCap ECs in different aging index groups (Fig. 8f). Costimulatory molecules, including *Cd80* and *Cd86*, peaked at aging index 5 but declined at index 6, while coinhibitory molecules progressively increased as the aging index rose, reaching their highest expression at index 6. These findings raise the possibility that alveolar capillary ECs may initially promote T cell activation during early aging, but shift toward suppression of T cell responses at more advanced aging states.

Given that IFNγ is a major inducer of MHC-related gene expression^52^, we next investigated whether the age-associated increase in MHC gene expression is mediated by IFNγ signaling. To achieve this, we treated aged mice with an IFNγ-neutralizing antibody and examined the expressions of MHC-related genes in their ECs (Fig. 8g). Expressions of *Cd74* and *H2-Ab*1, but not *B2m*, were significantly reduced following IFNγ neutralization (Fig. 8h). Flow cytometric and immunofluorescence analyses also revealed treatment with the IFNγ-neutralizing antibody to decrease cell-surface expression of MHC-II, but not MHC-I, in pulmonary and renal ECs (Fig. 8i,j). These findings suggest that IFNγ signaling contributes to the age-associated induction of MHC-II-related genes and cell-surface MHC-II expression in ECs.

Recent studies have shown that the gut microbiota changes compositionally with aging, contributing to age-related functional decline and disease development^53–56^.

Moreover, commensal microorganisms have been demonstrated to modulate IFN signaling and immune responses, including the induction of IFNγ-producing CD8 T cells by a specific gut bacterial consortium^57–59^. We therefore investigated whether age-associated MHC expression in ECs requires commensal microbial signals using aged germ-free mice. Cell-surface expressions of both MHC-I and MHC-II in pulmonary and renal ECs differed minimally between aged specific pathogen free mice and aged germ-free mice (Fig. 8k). Consistent with this observation, the age-associated expressions of *B2m*, *Cd74*, and *H2-Ab1* in ECs from the lungs were not affected by germ-free conditions (Fig. 8l). These results suggest age-associated induction of MHC-related genes in ECs to largely be independent of commensal microbial signals.

These findings, taken together, indicate that aging broadly enhances endothelial antigen presentation programs across multiple organs, with IFNγ signaling contributing to age-associated MHC-II induction independently of commensal microbiota, and suggest that alveolar capillary ECs may acquire an increasingly coinhibitory antigen presentation phenotype with advancing age.

## Discussion

In the present study, we performed scRNA-seq of ECs from five organs throughout the lifespan, and found that aging progressively causes erosion of organ-specific endothelial programs while inducing shared interferon-responsive and antigen-presentation programs across organs. Although vascular subtype and conserved capillary subset identities were largely preserved, organ-specific transcriptional programs were broadly attenuated with aging, indicating that gradual loss of organ-specific endothelial identity is a fundamental feature of endothelial aging. Importantly, these alterations were associated with deterioration of organ-specific endothelial functions, suggesting that age-related EC alterations compromise tissue homeostasis and resilience in processes impacting most, perhaps all, organs. Furthermore, we established single-cell aging indices for alveolar capillary ECs in mice and humans that quantitatively captured endothelial aging and revealed marked heterogeneity among ECs of the same chronological age. Notably, alveolar capillary ECs exhibited progressive functional decline before reaching a senescent-like state, suggesting loss of organ-specific endothelial programs to precede overt cellular senescence during aging.

In all organs examined, aging simultaneously induced attenuation of organ-specific endothelial programs and upregulation of interferon-responsive programs. Furthermore, aging indices of alveolar capillary ECs were largely determined by IFN-responsive genes, and increasing aging indices were associated with progressive loss of organ-specific endothelial programs. These observations raise the possibility that IFN signaling contributes to erosion of organ-specific endothelial programs during aging.

Previous studies have, in fact, identified chronic activation of IFN signaling as a driver of age-associated functional decline in multiple cell types^60–63^. Conversely, senescent cells are also known to induce chronic inflammation and tissue dysfunction through expression of IFN-stimulated genes^14,64,65^. Therefore, whether IFN signaling acts as a driver or is a consequence of endothelial aging remains an unresolved issue. IFN signaling may contribute to the loss of organ-specific endothelial programs, but it may also represent a secondary response to endothelial dysfunction and senescence.

Brain capillary ECs exhibited concurrent maladaptive and compensatory transcriptional responses during aging, resulting in both detrimental and protective alterations in BBB-associated gene programs. These opposing responses may partially preserve BBB integrity and homeostasis, thereby limiting age-related erosion of organ-specific endothelial functions relative to other organs. Nevertheless, further investigation is warranted to validate these findings and their functional implications.

IFNγ signaling, but not commensal microbial signals, regulates age-associated MHC-II expressions in pulmonary and renal ECs. As effector memory and activated CD4^+^ T cells increase in the lungs with aging while naive CD4^+^ T cells decline^66^, aged alveolar capillary ECs may stimulate CD4^+^ T cells via MHC-II-mediated antigen presentation. However, the expressions of coinhibitory molecules were upregulated in alveolar capillary ECs as the aging index rose. These findings raise the possibility that alveolar capillary ECs may initially promote T cell activation during early aging, but shift toward suppression of T cell responses at later stage of aging. Notably, this shift is consistent with recent studies demonstrating that senescent cells are capable of evading immune surveillance through upregulation of PD-L1^67–69^. Nevertheless, whether MHC-II-expressing aged ECs contribute to suppression of T-cell responses and/or altered T-cell regulation during aging awaits further detailed investigations.

We can reasonably speculate that stress-associated EC activation drives progressive functional deterioration and subsequent transition toward senescence in alveolar capillary ECs. Our aging index analysis revealed that intermediate State 3 ECs expressed not only the genes known to be associated with endothelial activation but also those involved in cell cycle arrest, stress responses, and DNA damage. These cells subsequently became IFN-responsive State 4 cells and ultimately senescent-like State 5 ECs. This transcriptional profile of State 3 ECs resembles the early stages of stress-induced senescence. Persistent cellular activation induced by oncogenic stress is known to trigger cellular senescence^70,71^. In addition, a recent study showed cyclin D1 and Cdk6, which are widely recognized as classic cell-cycle promoters, to also induce DNA damage and inflammatory responses during aging and senescence^72^. Therefore, stress-associated endothelial activation might represent a key intermediate stage linking age-related functional deterioration to IFN activation and cellular senescence, although further studies are needed to establish its causal role in endothelial aging.

Alveolar capillary ECs exhibited progressive functional decline before reaching fully senescent states, suggesting that the deterioration of endothelial functions precedes overt cellular senescence during aging. It is now well established that senescent cells exhibit the SASP, thereby contributing to chronic inflammation, tissue dysfunction, and age-related diseases via the release of inflammatory cytokines and chemokines^73,74^. Our findings extend this concept by implicating deterioration of organ-specific endothelial functions as a factor emerging prior to overt cellular senescence and thereby contributing to age-associated tissue dysfunction before the accumulation of fully senescent ECs. Thus, our results provide a framework for elucidating the processes underlying endothelial aging as a progressive loss of endothelial specialization. Our findings also highlight pre-senescent dysfunctional EC states as potential therapeutic targets.

## Methods

### Mice

Aged mice used in this study were primarily provided by Kobe University through the support of the Japan Agency for Medical Research and Development (AMED). Young mice and some aged mice were obtained from The Jackson Laboratory Japan (Yokohama, Japan), CLEA Japan (Tokyo, Japan), SLC Inc. (Hamamatsu, Japan), and Sankyo Labo Service Corporation (Tokyo, Japan). Most mice had a C57BL/6J background, except germ-free mice, which had a C57BL/6N background. Both male and female mice were used, with males comprising the majority. *Cd74*-P2A-EGFP knock-in mice were generated using the CRISPR/Cas9 system. A synthesized P2A-EGFP cassette (Eurofins Genomics) was inserted in place of the stop codon in exon 9 of the *Cd74* gene to enable endogenous expression of EGFP in *Cd74*-expressing cells. For IFN-γ neutralization, 2-year-old mice were intraperitoneally administered 500 μg of anti-mouse IFN-γ neutralizing antibody (clone XMG1.2; Leinco Technologies, #I-1119) or isotype control IgG (Leinco Technologies, #I-1195) every 3 days for a total of three doses. All animal experiments were approved by the Institutional Animal Care and Use Committee of Nippon Medical School and conducted in accordance with institutional guidelines.

### Single-cell RNA sequencing

#### Isolation of pulmonary endothelial cells

Female mice at 12, 29, 54, 65, 75, and 106 weeks of age (n = 2, 1, 2, 2, 2, and 3, respectively) were used for this analysis. Pulmonary ECs were isolated as previously described^75,76^. Mice were anesthetized with a mixture of medetomidine, midazolam, and butorphanol, and perfused with ice-cold phosphate buffered saline (PBS) through the right ventricle, followed by tracheal cannulation. Lungs were inflated with 1 ml of Dispase solution (25 U/ml; Thermo Fisher Scientific, #17105041), harvested, and minced in 5 ml of digestion buffer containing low-glucose DMEM, 100 μg/ml Liberase TM (Roche, #05401119001), 0.1 mg/ml DNase I (Sigma, #DN25), and 25 mM HEPES (Gibco, #15630-106). Tissues were incubated at 37°C for 30 min with mechanical trituration every 5 min. The reaction was quenched with PBS containing 2% fetal bovine serum (FBS), filtered through a 100 μm mesh, and centrifuged at 330 × g for 5 min. Erythrocytes were removed by incubation in 5 ml of 1× Red Blood Cell Lysing Buffer (pluriSelect, #60-00050-11) for 4 min. After neutralization and filtration through a 40 μm mesh (Greiner Bio-One, #542140 or #542040), cells were incubated with an antibody cocktail for 30 min on ice, including FITC-conjugated anti-CD45 (1:400; BioLegend, #103107), anti-TER119 (1:400; BioLegend, #116205), anti-CD326/EpCAM (1:400; BioLegend, #118207), and anti-CD41 (1:400; BioLegend, #133903), and PE-conjugated anti-CD31 (1:200; BioLegend, #102507). SYTOX Blue (1:5,000; Thermo Fisher Scientific, #S34857) was added for dead cell exclusion.

CD31⁺CD45⁻TER119⁻EpCAM⁻CD41⁻ ECs were sorted using a FACSAria Fusion cell sorter (BD Biosciences).

### Isolation of renal endothelial cells

Male mice at 11 and 106 weeks of age (n = 2 per group) were transcardially perfused with 20 ml of ice-cold D-PBS (NACALAI TESQUE, #14249-24). Renal ECs were isolated as previously described^22^. Kidneys were harvested, decapsulated, and minced into 1 mm³ pieces. Tissues were dissociated in 2.5 ml of digestion buffer (100 μg/ml Liberase TM, 0.1 mg/ml DNase I, and 25 mM HEPES in high-glucose DMEM) via three to four cycles of 5 min incubation at 37°C with manual trituration. The undigested fraction containing glomeruli was recovered by passage through a 40 μm mesh, centrifuged at 330 × g for 5 min, and further dissociated in 0.1% trypsin (Cosmo Bio, #16205004) and 0.05 mg/ml DNase I for 25–30 min at 37°C. Pooled single-cell fractions were treated with 1× Red Blood Cell Lysing Buffer for 4 min at room temperature. Cells were stained for 45 min at 4°C with PE-conjugated anti-CD31 (1:200), PE-conjugated anti-Endomucin (1:100; Invitrogen, #12-5851-82), and Alexa Fluor 647-conjugated Isolectin GS-IB4 (1:500) for positive selection, and FITC-conjugated anti-TER119, anti-CD45, and anti-CD41 (all 1:400) for negative selection. SYTOX Blue (1:5,000) was added for viability staining.

CD31⁺Endomucin⁺IB4⁺CD45⁻TER119⁻CD41⁻ ECs were sorted using a FACSAria Fusion cell sorter.

### Isolation of liver endothelial cells

Liver ECs were isolated as previously described^77^. Male mice at 12 and 106 weeks of age (n = 1 and 2, respectively) were used. Livers were perfused through the intrahepatic inferior vena cava using a 24-gauge needle (Terumo, #SR-OT2419C). Clearing buffer containing 1× Hanks’ Balanced Salt Solution (HBSS; Wako, #082-09865), 0.5 mM EGTA, 0.5 mM EDTA, 20 mM HEPES, and 1 mM flavopiridol hydrochloride hydrate (Sigma, #F3055) was perfused at 5 ml/min for 3 min, followed by 40 ml of liver digestion buffer B containing 0.1 mg/ml Liberase TM and 1 μM DNase I (Worthington, #LS210039) in liver digestion buffer A. Liver digestion buffer A consisted of 20 mM HEPES and 1 μM flavopiridol in 1× HBSS containing calcium and magnesium. The liver was minced and incubated in 20 ml of digestion buffer B at 37°C for 20 min on a rotator. The suspension was filtered through a 100 μm mesh and centrifuged at 50 × g for 3 min twice to isolate non-parenchymal cells (NPCs). NPCs were purified using a 20% isotonic Percoll (GE Healthcare, #17-0891-01) gradient followed by OptiPrep (Cosmo Bio, #AXS-1114542) density gradient centrifugation at 1,400 × g for 25 min with lowest acceleration and no brake. After red blood cell lysis for 5 min on ice, cells were washed with PBS. NPCs were incubated with Fc blocker (1:500; BioLegend, #101309) for 10 min, centrifuged at 300 × g for 5 min, and stained with PE-conjugated anti-CD31 (1:200), and APC-conjugated anti-CD45 and anti-CD41 (1:200; BioLegend, #133903) for 30 min at 4°C. SYTOX Blue (1:2,000) was added for viability staining. CD31⁺CD45⁻CD41⁻ LSECs were sorted using a FACSAria Fusion cell sorter.

### Isolation of cardiac endothelial cells

Cardiac ECs were isolated as previously described^78^. Male mice at 3 and 24 months of age (n = 3 per group) were transcardially perfused with 20 ml of D-PBS per body at a flow rate of 3–5 ml/min. Hearts were minced and dissociated in buffer containing Knockout DMEM (Gibco, #10829018), MEM-NEAA (Gibco, #11140050), 1 mM sodium pyruvate (Gibco, #11360070), 4 μl/ml ECGS/H (PromoCell, #C-30120), 1 mg/ml Collagenase II (Gibco, #17101-015), 2.5 mg/ml Collagenase IV (Worthington, #LS004188), and 37.5 μg/ml DNase I for 25 min at 37°C, followed by mechanical dissociation using a gentleMACS Dissociator (Miltenyi Biotec, #130-093-235). ECs were enriched by magnetic-activated cell sorting (MACS) using CD31 MicroBeads (Miltenyi Biotec, #130-097-418). Enriched cells were stained with PE-conjugated anti-CD31, FITC-conjugated anti-CD45, anti-TER119, and anti-CD41 (all 1:200), and APC-conjugated anti-CD140a (1:200; eBioscience, #17-1401-81) for 30 min on ice. SYTOX Blue (1:2,000) was added for viability staining. CD31⁺CD45⁻TER119⁻CD41⁻CD140a⁻ ECs were sorted using a FACSAria Fusion cell sorter.

### Isolation of brain endothelial cells

Male mice at 3, 12, and 24 months of age (n = 2, 1, and 2, respectively) were transcardially perfused with 40 ml of heparinized PBS (1 U/ml heparin; Terumo, #SV-25DLK). Brain tissues were dissociated using the Neural Tissue Dissociation Kit– Postnatal Neurons (Miltenyi Biotec, #130-094-802). Tissues were incubated in Enzyme Mix 1 for 15 min at 37°C, mechanically dissociated using 90°-cut needles (TOP, #2865-10), and further incubated with Enzyme Mix 2 for 10 min. Myelin was removed using Myelin Removal Beads II (Miltenyi Biotec, #130-096-733) for 15 min at 4°C. ECs were enriched by MACS using CD31 MicroBeads. Cells were stained with FITC-conjugated anti-CD45 and anti-CD41, and PE-conjugated anti-CD31 (all 1:200) for 30 min on ice. SYTOX Blue (1:2,000) was added for viability staining. CD31⁺CD45⁻CD41⁻CD140a⁻ ECs were sorted using a FACSAria Fusion cell sorter.

### Single-cell suspension preparation and library construction

Following sorting, ECs were centrifuged at 300 × g for 8 min at 4°C. The supernatant was aspirated, and the cell pellet was resuspended in 1.0 ml of FACS buffer (2% FBS in D-PBS). The suspension was transferred to a 1.5 ml tube, adjusted to 1.3 ml, and washed twice by centrifugation at 500 × g for 4 min at 4°C. After the final wash, the supernatant was carefully removed, leaving approximately 40 μl, and cells were resuspended in FACS buffer to a concentration of 1,000 cells/μl. The suspension was filtered through a Flowmi tip strainer (Bel-Art Products, #136800040) to remove aggregates. Cell concentration and viability were assessed by Trypan Blue (0.4%; Gibco, #15250-061) exclusion using a hemocytometer or Countess 3 (Thermo Fisher Scientific), with counts performed in duplicate.

Single-cell RNA-seq libraries were constructed using the Chromium Next GEM Single Cell 3’ GEM, Library & Gel Bead Kit v3.1 (10x Genomics, #PN-1000128) and Chromium Next GEM Chip G Single Cell Kit (10x Genomics, #PN-1000127) on a Chromium Controller (10x Genomics) according to the manufacturer’s instructions.

Library quality was assessed using a 2100 Bioanalyzer (Agilent Technologies) with the High Sensitivity DNA Kit (Agilent, #5067-4626) or a Qubit fluorometer (Thermo Fisher Scientific). Libraries were converted to single-stranded circular DNA compatible with the DNBSEQ-G400RS platform (MGI Tech) using the MGIEasy Universal Library Conversion Kit App-A (MGI Tech, #1000004155) according to the manufacturer’s instructions. Sequencing reads were processed using Cell Ranger Single Cell Software Suite (v7.1.0) with a custom annotation of the mouse genome assembly (mm10) to perform barcode processing and gene counting with default parameters.

### Data preprocessing and annotation

Sequencing metrics, including mean sequencing depth, percentage of uniquely mapped reads, median number of genes per cell, and total cell counts after filtering, are summarized in Table S1. Unless otherwise noted, all scRNA-seq analyses, including clustering and differential expression, were performed using the Seurat R package (v5.2.1)^79^. Low-quality cells and minimally-expressed genes were excluded using dataset-specific filtering thresholds detailed in Table S2. Filtered datasets were merged, and standard preprocessing was applied, including NormalizeData(), FindVariableFeatures(), ScaleData(), RunPCA(), FindNeighbors(), FindClusters(), and RunUMAP() prior to batch correction. Data integration was performed using canonical correlation analysis (CCA), followed by JoinLayers(), FindNeighbors(), FindClusters(), and RunUMAP() to visualize integrated clusters. Doublets were identified based on nFeature_RNA and nCount_RNA thresholds.

Cluster annotation was performed using a three-level strategy based on marker gene expression (Table S3). At level 1, EC clusters within tissues from each organ were identified to exclude non-endothelial cells. At level 2, EC clusters were sub-clustered and annotated into vascular subtypes with reference to previously published studies (lungs^17,80,81^, kidneys^22,82^, liver^8,24^, heart^83–85^, brain^10,86,87^ (Table S4). At level 3, capillary populations identified at level 2 were isolated and further sub-clustered based on conserved marker genes within each tissue sample (Figures S3C and Table S5).

### Bioinformatics Analysis

#### Organ heterogeneities (related to Figure1)

DEGs were defined as genes with an absolute log2 fold-change (|log2FC|) > 0.5, identified using FindAllMarkers() for comparisons between clusters in young mice (restricted to positive markers, only.pos = TRUE) and FindMarkers() for comparisons between age groups (including both upregulated and downregulated genes). For heatmap visualization, the top 100 or 300 DEGs were selected, ranked by adjusted p-value and then by descending log2 fold-change. For Gene Ontology (GO) analysis, the top 100 DEGs were selected after excluding ribosomal protein genes and sex-specific genes (Table S6), and analyzed using the clusterProfiler R package (v4.10.1)^88^. Module scores were calculated based on the top 300 tissue-specific genes identified in young tissues and compared between young and aged tissues using violin plots or feature plots. For the top 300 age-related DEGs (downregulated and upregulated separately), semantic similarity between GO Biological Process terms was evaluated using the GOSemSim R package (v2.28.1)^89^ with Wang’s method and the best-match average (BMA) strategy for score aggregation, and visualized as heatmaps using the pheatmap (v1.0.13)^90^ and ggplot2 (v3.5.2)^91^ R packages.

### Analysis of Vascular Subtype Heterogeneity (related to Figure 2)

To identify robust marker genes for vascular subtypes (artery, arteriole, capillary, venule, and vein) across vascular beds, ECs were isolated from each young tissue, and subtype-specific markers were identified using FindAllMarkers() with a log2 fold-change threshold of >0.25 for capillaries and >0.5 for all other subtypes (min.pct = 0.25, only.pos = TRUE). Tissue-specific marker lists were merged to extract conserved markers consistently expressed in each vascular subtype across organs. Module scores for conserved markers and tissue-specific markers within each of the 25 groups (five vascular subtypes across five tissues) were calculated as described above and z-scores were normalized to enable direct comparison between vascular subtype and tissue-specific signatures. Tissue heterogeneity among the 25 groups was evaluated by principal component analysis (PCA) using the top 2,500 variable genes identified by FindVariableFeatures() and visualized using the plotly R package (v4.11.0) ^92^. Statistical analyses and additional visualizations were performed using GraphPad Prism 10 (GraphPad Software).

### Aging-based pseudotime analysis (related to Figures 3, 5, and 6)

To characterize transcriptomic dynamics within specific capillary subsets (lungs: gCap-IFNg, gCap-AP-1, and gCap-Signal; heart: all subtypes; liver: Cap-Artery-like, Cap-Vein-like, Cap-Signal, and Cap-AP-1), the tradeSeq R package (v1.16.1)^93^ was used.

Module scores for aging index-defining genes were calculated for individual cells using AddModuleScore() in Seurat and used to construct an aging-based pseudotime. Genes exhibiting statistically significant temporal changes along this pseudotime were identified using the associationTest() function in tradeSeq, and the peak expression time for each significant gene was determined to order genes along the pseudotime trajectory. Pseudotime was divided into 100 bins, and average expressions of aging index-defining genes and functional genes within each bin were calculated using AverageExpression() and visualized as heatmaps using the pheatmap R package, illustrating the stepwise decline of gene expression associated with the aging signature. For lungs, line graphs were generated from the average expressions of genes within the gCap-IFN, gCap-AP-1, and gCap-Signal subpopulations, normalized to the value at 12 weeks.

### Analysis of the Integrated Human Lung Cell Atlas (HLCA) Data

The HLCA dataset was obtained in h5ad format from Figshare^21^. Cells annotated as “capillary endothelial cell” within the “lung parenchyma” were subsetted from the original HLCA object (28,024 features across 20,210 cells). The corresponding gene expression matrix and metadata were reconstructed into a Seurat object (v5.2.1) and normalized using NormalizeData(). To investigate natural aging, “healthy” donors with no history of smoking (as defined in the “lung_condition” metadata) were categorized into three age groups: young (<40 years, n = 8), middle-aged (40–59 years, n = 8), and aged (≥60 years, n = 7). Expression heatmaps were generated based on average gene expression per donor using AverageExpression(). An aging index for human cells was calculated by determining the module score for aging index-defining genes (Table S7) using AddModuleScore(). To evaluate the effect of smoking on the aging index, “healthy” donors in each age group were further stratified by smoking status (young never-smokers, n = 8; young former smokers, n = 4; young current smokers, n = 6; aged never-smokers, n = 7; aged current smokers, n = 1).

### RNAscope in situ hybridization

RNAscope in situ hybridization was performed using the RNAscope Multiplex Fluorescent Reagent Kit v2 (Advanced Cell Diagnostics, Newark, CA) according to the manufacturer’s instructions, with modifications for TSA-based detection. Probes were purchased from Advanced Cell Diagnostics: *Pecam1* (#316721), *Sftpc* (#314101), *Scgb1a1* (#420351), *Ptprm* (#579201), *Ifit3* (#508251), *Alb* (#437691), *Tnnt2* (#880391), *Miox* (#1086421), and *Cacna1e* (#449211). Sections were baked at 60°C for 30 min, deparaffinized, and treated with hydrogen peroxide for 10 min at room temperature. For target retrieval, slides were boiled in Target Retrieval Buffer at 98– 102°C for 1 min, dehydrated in 100% ethanol, and treated with Protease III at 40°C for 30 min. Hybridization was performed with a pooled probe cocktail including C1 (*Pecam1*), C2 (*Ptprm*, *Scgb1a1*, *Miox*, *Tnnt2*, *Alb*), and C3 (*Ifit3*, *Sftpc*, *Cacna1e*) for 2 h at 40°C. Signal amplification was performed by sequential hybridization with AMP1 and AMP2 (30 min each at 40°C) and AMP3 (15 min at 40°C). Fluorescence detection was performed sequentially: C1 with HRP-C1 (15 min) followed by TSA-Vivid 520 (30 min); C2 with HRP-C2 (15 min) followed by TSA-Vivid 570 (30 min); and C3 with HRP-C3 (15 min) followed by TSA-Vivid 650 (30 min). HRP blocker was applied for 15 min at 40°C between each fluorophore to prevent signal cross-talk. Nuclei were counterstained with DAPI for 30 s at room temperature. Slides were mounted, and images were acquired using an AX-R confocal microscope (Nikon).

### Immunofluorescence Staining

#### Lungs from *Cd74*-P2A-EGFP knock-in, naturally aged, and anti-IFN-γ-treated mice

Mice were deeply anesthetized with a mixed anesthetic (0.3 mg/kg medetomidine, 4.0 mg/kg midazolam, and 5.0 mg/kg butorphanol; 100 μl per 10 g body weight). Airways were inflated via the trachea with 1–3 ml of 1% low-melting agarose (NACALAI TESQUE, #01651-92) in PBS (pre-warmed to 37°C) until the lung lobes were fully expanded, then immediately cooled on ice for 2 min to solidify the agarose. Lungs were excised and post-fixed in 4% paraformaldehyde (NACALAI TESQUE, #09154-85) at 4°C for 60 min to harden the tissue surface. Lung lobes were separated and sectioned at 200–300 μm using a vibratome (Leica VT1200) in pre-chilled PBS, and thick sections were further fixed in 4% paraformaldehyde at 4°C for 60 min on ice.

Sections were blocked and permeabilized with 5% normal goat serum or normal donkey serum (Jackson ImmunoResearch, #017-000121) in 0.25% Triton X-100/PBS containing sodium azide at 4°C for 30 min. Sections were incubated overnight at 4°C with primary antibodies diluted in blocking buffer: anti-GFP (1:1000; Thermo Fisher Scientific, #A-11122), anti-CD31 goat (1:400; R&D Systems, #AF3628), anti-CD31 rat (1:200; BD Biosciences, #550274), anti-VE-cadherin (1:400; R&D Systems, #AF1002), and APC-conjugated anti-CD45 (I3/2.3) (1:200; BioLegend, #147707). After washing with PBS containing sodium azide (15 min, twice), sections were incubated overnight at 4°C with secondary antibodies: Alexa Fluor 488-conjugated donkey anti-rabbit (Jackson ImmunoResearch, #705-545-147), Alexa Fluor 488-conjugated donkey anti-rat (Jackson ImmunoResearch, #712-545-153), Cy3-conjugated anti-goat (Jackson ImmunoResearch, #705-165-003), and Cy3-conjugated anti-rabbit (Jackson ImmunoResearch, #711-165-152), together with DAPI (1 μM; Invitrogen, #D1306). After a final wash, the sections were mounted with mounting medium (Abcam, #ab104135) for imaging.

### Immunofluorescence staining of cardiac tissue

Snap-frozen hearts were cryosectioned at 10 μm and mounted on CREST-coated glass slides (Matsunami). Sections were equilibrated from −80°C to room temperature for 30–60 min, fixed with 4% PFA in PBS for 15 min at room temperature, and permeabilized with 0.25% Triton X-100 in PBS for 10 min. After blocking with 3% normal donkey serum in 0.1% Triton X-100/PBS for 1 h at room temperature, sections were incubated overnight at 4°C with primary antibodies: goat anti-CD31 (1:400; R&D Systems, #AF3628) and rabbit anti-Ki67 (1:300; Abcam, #ab15580). After washing with PBS, sections were incubated with secondary antibodies—Alexa Fluor 488-conjugated donkey anti-goat (1:400; Jackson ImmunoResearch, #705-545-147) and Cy3-conjugated donkey anti-rabbit (1:400; Jackson ImmunoResearch, #711-165-152)— for 1 h at room temperature.

To enable sequential staining with another rabbit-derived antibody, sections were re-fixed with 4% paraformaldehyde for 15 min at 4°C and blocked with 5% normal rabbit serum (Fujifilm, #140-06571) in 0.1% Triton X-100/PBS for 30 min at 4°C. Sections were then incubated for 4–5 h at 4°C with Alexa Fluor 647-conjugated rabbit anti-ERG (1:300; Abcam, #ab196149) and DAPI (1.5 µM; Thermo Fisher Scientific) diluted in the same blocking buffer. Sections were rinsed, treated with 0.45% NaCl, and mounted using ProLong Glass Antifade Mountant (Thermo Fisher Scientific, #P36984).

### Immunofluorescence staining of liver tissue

Thick liver sections (200 μm) were prepared using a vibratome and stored in 50% glycerol at −30°C until use. Sections were blocked with 5% normal serum and 0.25% Triton X-100 in PBS containing sodium azide for 30 min at 4°C. Primary antibody incubation was performed overnight at 4°C using one of the following combinations: rabbit anti-c-Jun (1:200; Cell Signaling Technology, #9165) and goat anti-CD31 (1:200; R&D Systems, #AF3628) for c-Jun staining, or rabbit anti-CD34 (1:200; Abcam, #ab81289) and rat anti-CD31 (1:200; BD Biosciences, #550274) for CD34 staining.

After washing three times with PBS containing sodium azide (30 min each) at 4°C, sections were incubated overnight at 4°C with appropriate fluorophore-conjugated secondary antibodies (1:500) and 3 μM DAPI. For ERG labeling, sections were post-fixed with 4% PFA for 2–3 h at 4°C, blocked with 5% normal rabbit serum, and incubated overnight at 4°C with Alexa Fluor 647-conjugated anti-ERG (1:500). Sections were washed three times and mounted for confocal imaging.

### Immunofluorescence staining of kidney tissue

Kidneys were harvested from mice transcardially perfused with heparinized PBS and fixed with 4% paraformaldehyde in PBS for 2–3 h at 4°C. After fixation and removal of the renal capsule, one side of each kidney was trimmed using a surgical knife. Thick sections (200–300 μm) were prepared using a vibratome and further fixed with 4% paraformaldehyde in PBS for 2–3 h at 4°C. Sections were blocked for 30 min at 4°C as described above and incubated overnight at 4°C with primary antibodies: rabbit anti-c-Jun (1:200; Cell Signaling Technology, #9165) and goat anti-CD31 (1:200; R&D Systems, #AF3628). After washing with PBS containing sodium azide (30 min each) at 4°C, appropriate secondary antibodies were applied overnight at 4°C together with 1 μM DAPI. For ERG labeling, additional steps were performed as described in “Immunofluorescence staining of liver tissue.”

### Picrosirius red staining of cardiac tissue

Mice were intracardially perfused with 20–25 ml of 1× PBS per animal. Hearts were dissected and fixed in 4% PFA overnight at 4°C, then embedded in paraffin and sectioned at 4 µm thickness. Sections were deparaffinized, rehydrated, and stained with Weigert’s iron hematoxylin (Muto Pure Chemicals, #40342 and #40352) for 10 min, followed by differentiation. Sections were then stained with 0.03% picrosirius red solution (Muto Pure Chemicals, #33061 and #40362) for 12 min at room temperature. Images were acquired using a VS200 slide scanner (Evident) with a 40× objective lens. Image analysis was performed using QuPath software (version 0.7.0, University of Edinburgh). Collagen-positive areas were identified by training a pixel classifier using the “Train Pixel Classifier” function to detect red-stained pixels. Interstitial collagen area was calculated by subtracting the epicardium, endocardium, and large vessel areas from the total tissue area.

### Scavenging activity assay in LSECs

Primary liver ECs were isolated using a combination of enzymatic digestion and magnetic-activated cell sorting (MACS). Livers were excised, minced in PBS, and digested in DMEM containing 1 mg/ml Liberase (Roche/Sigma-Aldrich) for 30 min at 37°C. The cell suspension was filtered through a cell strainer and diluted to 50 ml with FACS buffer (PBS containing 2% FBS and 2 mM EDTA). To remove parenchymal hepatocytes, the suspension was centrifuged at 50 × g for 2 min at 4°C, and the supernatant containing NPCs was collected; this low-speed centrifugation step was repeated to maximize hepatocyte removal. NPCs were recovered by centrifugation at 450 × g for 5 min at 4°C. Red blood cells were lysed for 3 min at 4°C, followed by being washed with PBS. The NPC pellet was resuspended in FACS buffer and incubated with CD45 MicroBeads (Miltenyi Biotec) at a 1:9 ratio for 15 min at 4°C to deplete or enrich the leukocyte fraction as required, followed by separation using an LS column (Miltenyi Biotec).

The EC-enriched fraction was seeded at 3 × 10⁵ cells per dish in endothelial cell growth medium consisting of high-glucose DMEM supplemented with 20% FBS, 50 μg/ml Endothelial Cell Growth Supplement (Corning), 100 μg/ml heparin (Sigma-Aldrich), and penicillin/streptomycin. Medium was replaced 20 h after seeding to remove non-adherent cells. At 24 h post-seeding, cells were incubated with 10 μg/ml Alexa Fluor 488-conjugated OVA (Thermo Fisher Scientific, #O34781) in serum-free DMEM for 2 h at 37°C to allow endocytosis. Cells were washed with D-PBS, fixed with 4% paraformaldehyde for 15 min at room temperature, and permeabilized with 0.1% Triton X-100 in PBS containing 2.5% donor serum for 30 min. Cells were stained with goat anti-CD31 (1:500) overnight at 4°C, followed by Cy3-conjugated anti-goat secondary antibody (1:1000) for 1.5 h at room temperature, and then incubated with Alexa Fluor 647-conjugated anti-ERG (1:400) and DAPI for 2 h at room temperature. Images were acquired using a Nikon AX-R confocal microscope.

### Vascular permeability assay in the lungs

To induce acute lung injury, mice were anesthetized by intraperitoneal injection of a mixed anesthetic (0.3 mg/kg medetomidine, 4.0 mg/kg midazolam, and 5.0 mg/kg butorphanol; 100 μl per 10 g body weight). Lipopolysaccharide (LPS; 4 mg/kg; Sigma-Aldrich, #L2880) diluted in saline to a total volume of 50 μl was administered intranasally in small increments to both nares using a pipette with a modified tip to ensure complete inhalation. Mice were maintained in an upright position for 5 min after administration, and anesthesia was reversed by intraperitoneal injection of atipamezole (100 μl per 10 g body weight).

Vascular permeability was assessed 24 h after LPS administration using the Evans Blue extravasation assay. Mice were anesthetized, and 1% Evans Blue (Fluka, #46160) in PBS (50 mg/kg) was injected into the retro-orbital venous sinus using a 29-gauge needle; successful systemic distribution was confirmed by blue discoloration of the extremities. Thirty minutes after injection, blood was collected from the tail vein and serum was separated using a serum-separating tube (Tochitube, Sato Kasei Kogyosho) according to the manufacturer’s instructions, then stored at −80°C. Mice were then euthanized and transcardially perfused with approximately 30 ml of HBSS (Wako, #082-09865) through the right ventricle to remove intravascular dye. Lungs were harvested, washed in ice-cold HBSS, divided into anatomical sections, and dried at 50°C for 24 h to determine dry tissue weight. Evans Blue was extracted by incubating dried tissues in 500 μl of formamide at 50°C for 24 h, and extracts were centrifuged at 19,000 × g for 5 min at 4°C. Absorbance of the supernatant and serum (diluted 1:100 in formamide) was measured at 620 nm using a microplate reader. The Evans Blue concentration in each tissue was calculated from a standard curve, normalized to dry tissue weight, and expressed relative to the serum Evans Blue concentration (lung/serum ratio).

### Flow cytometry analysis

To analyze the fraction of MHC-I- and MHC-II-positive ECs, flow cytometry was performed on cells isolated from each organ as described above. Cells were resuspended in D-PBS (NACALAI TESQUE, #14249-24) containing 2% FBS (lungs and kidneys) or PBS containing 0.5% BSA and 2 mM EDTA (heart), and stained for 30 min at 4°C with the following fluorochrome-conjugated antibodies: PE-conjugated anti-CD31 (BioLegend, #102507; 1:6400 for lungs, 1:3200 for kidneys, 1:5000 for liver, 1:3000 for heart, 1:1000 for brain), APC-conjugated anti-CD45 (BioLegend, #147707; 1:3200 for lungs, 1:1600 for kidneys, 1:1000 for liver, heart, and brain), Alexa Fluor 488-conjugated anti-MHC-I (BD Bioscience #567696; 1:200 for lungs, kidneys, and liver, 1:150 for heart and brain), and Alexa Fluor 488-conjugated anti-MHC-II (Biolegend #107615; 1:200 for lungs, kidneys, liver, and brain, 1:150 for heart). After washing with the corresponding buffer, as described above, cells were incubated with SYTOX Blue (1:5000 for lungs and kidneys, 1:3000 for heart) for 2 min (lungs and kidneys) or 10 min (heart). Flow cytometry was performed using a FACSAria Fusion cell sorter (BD Biosciences).

### qPCR Analysis

Total RNA was extracted from FACS-sorted ECs using TRIzol RNA Isolation Reagent followed by the Direct-zol RNA MicroPrep Kit (Thermo Fisher Scientific, #15596026; Zymo Research, #R2061) or the ISOSPIN Cell & Tissue RNA Kit (NIPPON GENE, #314-08211). cDNA was synthesized from 50 ng of total RNA using SuperScript IV VILO Master Mix (Thermo Fisher Scientific, #11756050). Transcript levels were quantified using THUNDERBIRD Next SYBR qPCR Mix (TOYOBO, #QPX-201) on a CFX96 Real-Time PCR Detection System (Bio-Rad). The following primer sets were used: mouse *Gapdh*, 5’-AGGTCGGTGTGAACGGATTTG-3’ and 5’-TGTAGACCATGTAGTTGAGGTCA-3’; mouse *Actb*, 5’-GTGACGTTGACATCCGTAAAGA-3’ and 5’-GCCGGACTCATCGTACTCC-3’; mouse *Pecam1*, 5’-CCATGGAAGAAAGGGCTCATTGC-3’ and 5’-CGTAATGGCTGTTGGCTTCCA-3’; mouse *Cd74*, 5’-ATGAACAGTTGCCCATACTGGGC-3’ and 5’-GCTGGTACAGGAAGTAAGCAGTGG-3’; mouse *B2m*, 5’-CACTGACCGGCCTGTATGCTAT-3’ and 5’-TCAATGTGAGGCGGGTGGAACT-3’; mouse *H2-Ab1*, 5’-GACACGGTGTGCAGACACAACTA-3’ and 5’-CAGACCAGAGTGTTGTGGTGGT-3’; mouse *Aplnr*, 5’-ACAGCAGTAGTGCTGAGAAGTCAG-3’ and 5’-TTTCTCATGCATCTGCTCTCCTCC-3’; mouse *Ptprm*, 5’-TGGACAAGTGGCAGGAGGAATACA-3’ and 5’-GACATCCACAGTTCTCTGGTGTCG-3’.

### Quantitative image analysis

#### Quantification of EC proliferation in the heart

To quantify EC density and proliferation in the heart, tilescan z-stack images of the left ventricular wall and apex were acquired using an AX-R confocal microscope (Nikon), and maximum intensity z-projections were generated from the z-stacks. Endothelial nuclei were automatically segmented based on the ERG signal using Cellpose-SAM, custom-trained for this dataset as previously described^94^. Segmentation masks were imported into Fiji/ImageJ to define regions of interest (ROIs) for individual ERG⁺ nuclei and determine total EC counts. Proliferating ECs were defined as ERG-positive ROIs with mean Ki67 channel intensity exceeding 1,000 arbitrary units. Proliferation rates were calculated as the percentage of Ki67- and ERG-positive cells relative to total ERG-positive cells. EC density was calculated as the total number of ERG-positive cells per unit tissue area.

### Quantification of c-Jun expression in hepatic endothelial cells

c-Jun-positive ECs were quantified using three-dimensional image analysis in Imaris software (Bitplane). Endothelial nuclei were identified using the Spot detection function based on the ERG channel, and the total number of detected spots was defined as the total EC count. c-Jun-positive signals were similarly identified using the Spot function. To determine endothelial-specific c-Jun expression, c-Jun spots located within 2.2 μm of any ERG spot were extracted and quantified as c-Jun-positive ECs. The percentage of c-Jun-positive ECs relative to the total EC count was calculated as a measure of EC activation.

### Quantification of OVA internalization in hepatic endothelial cells

OVA-Alexa Fluor 488 intensity within ECs was quantified using Fiji/ImageJ.

Multichannel confocal images were split into single-channel images, and the CD31 channel was thresholded to generate a binary mask. ROIs corresponding to individual cell boundaries were defined using the Analyze Particles tool, and non-endothelial regions lacking ERG expression were manually excluded. Within each validated ROI, nuclei were counted, and integrated density of the OVA-Alexa Fluor 488 signal was measured. To account for multinucleated ECs or overlapping cell boundaries, mean fluorescence intensity per cell was calculated by dividing the integrated density of each ROI by its corresponding nuclear count, and averaged across cells for each sample.

### Quantification of VE-cadherin junctional profiles

Junctional VE-cadherin distribution and intensity were quantified by line-scan analysis using Fiji/ImageJ. Confocal z-stack images (15 μm thickness) were converted to maximum intensity z-projections, converted to 8-bit format, and standardized for brightness and contrast across all experimental groups before being saved as JPEG files. Line scans (30 pixels in length) were drawn perpendicular to cell–cell borders, centered on the peak intensity of the VE-cadherin signal, and fluorescence intensity at each pixel along the line was recorded to generate spatial intensity profiles.

In total, 121–252 lines were measured per mouse across four age groups: 12, 54, 79, and 107 weeks (n = 3 mice per group), with three independent images evaluated per animal.

## Data and code availability

✓ scRNA-seq data obtained in this study have been deposited in the DNA Data Bank of Japan (DDBJ) and are publicly available as of the date of publication. The accession numbers are DRA029182 and E-GEAD-1278.
✓ This study generated a publicly accessible web-based database for browsing the data presented in this paper, available at moea-fukulab.org as of the date of publication.
✓ This study reanalyzed publicly available human lung scRNA-seq data from the integrated Human Lung Cell Atlas (HLCA)^21^, obtained via the HCA Data Portal (https://data.humancellatlas.org/hca-bio-networks/lung/atlases/lung-v1-0), governed by the HCA Data Release Policy (https://www.humancellatlas.org/data-release-policy) and licensed under CC BY 4.0.
✓ The code and data supporting the findings of this study will be made publicly available on GitHub as of the date of publication.
✓ Any additional information required to reanalyze the data reported in this paper is available from the lead contact upon request.

## Acknowledgements

We thank Y. Taguchi (BD Biosciences) for supporting the experiments using FACSAriaTM Fusion. We also thank Y. Matsushita and K. Kato for excellent technical assistance. We thank the Advanced Animal Model Support Platform (AdAMS) for support with mouse model generation. This platform is supported by JSPS KAKENHI Grant Number JP22H04922. Computations were partially performed on the NIG supercomputer at ROIS National Institute of Genetics. Picrosirius red staining was supported by the Pathology Core Laboratory, The Institute of Medical Science, The University of Tokyo. We thank the Human Lung Cell Atlas (HLCA) consortium, part of the Human Cell Atlas Lung Biological Network (coordinated by P. Barbry, M. Nawijn, J. Rajagopal and N. Banovich), and the Human Cell Atlas Data Coordination Platform for making the integrated lung scRNA-seq data publicly available.

This work was supported by AMED under Grant Number JP23gm1710009 to S.F. and JP22gm6710002 to M.S.; by Grants-in-Aid for Scientific Research (B) to S.F. (21H02665), to H.W.-T. (26K02221), and to S.T. (23K24371, 26K02221), for Scientific Research (C) to H.W.-T. (23K06325), and for Exploratory Research to S.F. (23K18245), all from the Japan Society for the Promotion of Science (JSPS); and by the FOREST Program from the Japan Science and Technology Agency (JST) under Grant Numbers JPMJFR220T to H.W.-T. and JPMJFR210S to S.T. This work was also supported by the Research Support Project for Life Science and Drug Discovery (Basis for Supporting Innovative Drug Discovery and Life Science Research (BINDS)) from AMED under Grant Number JP25ama121055 (Support Number 5081). This work was also supported by grants from the Naito Foundation to S.F., from Daiichi Sankyo Foundation of Life Science to S.F. and H.W.-T., from The Uehara Memorial Foundation to S.F., from the Mochida Memorial Foundation to H.W.-T., and the Ichiro Kanehara Foundation for the Promotion of Medical Science and Medical Care to H.W.-T.

## Author contributions

H.W.-T. and S.F. conceptualized the study, designed the experiments, interpreted the data, and wrote the manuscript. H.W.-T. performed experiments on lung and kidney ECs, conducted cross-tissue analyses, and oversaw the overall data analysis and integration. T.I. performed experiments on heart and brain ECs with help from H.M., T.T., S.H., and Y.R., conducted the corresponding data analysis, and contributed to the overall data analysis. T.H. performed experiments on liver ECs with help from D.H., S.T., and M.S., and analyzed the corresponding data. H.I., K.Y., and M.H. supported the data analysis and database development. E.O.-N. performed lung injury experiments.

K.A. conducted RNAscope in situ hybridization with help from S.H., M.M., and H.I. K.Ar. generated the *Cd74*-P2A-EGFP mouse line with help from T.M. S.T. supported FACS analysis and was involved in discussions on MHC data. All authors discussed the results and commented on the manuscript.

## Declaration of generative AI and AI-assisted technologies

During the preparation of this work, the authors used ChatGPT and Claude to assist with language editing, manuscript preparation, and supporting the implementation of established computational methods in analysis code. The authors reviewed and edited all AI-generated content as needed and take full responsibility for the content of the published article.

