## Supplementary Information for "Age-associated erosion of organ-specific endothelial programs compromises tissue function and resilience"

### Extended Figures

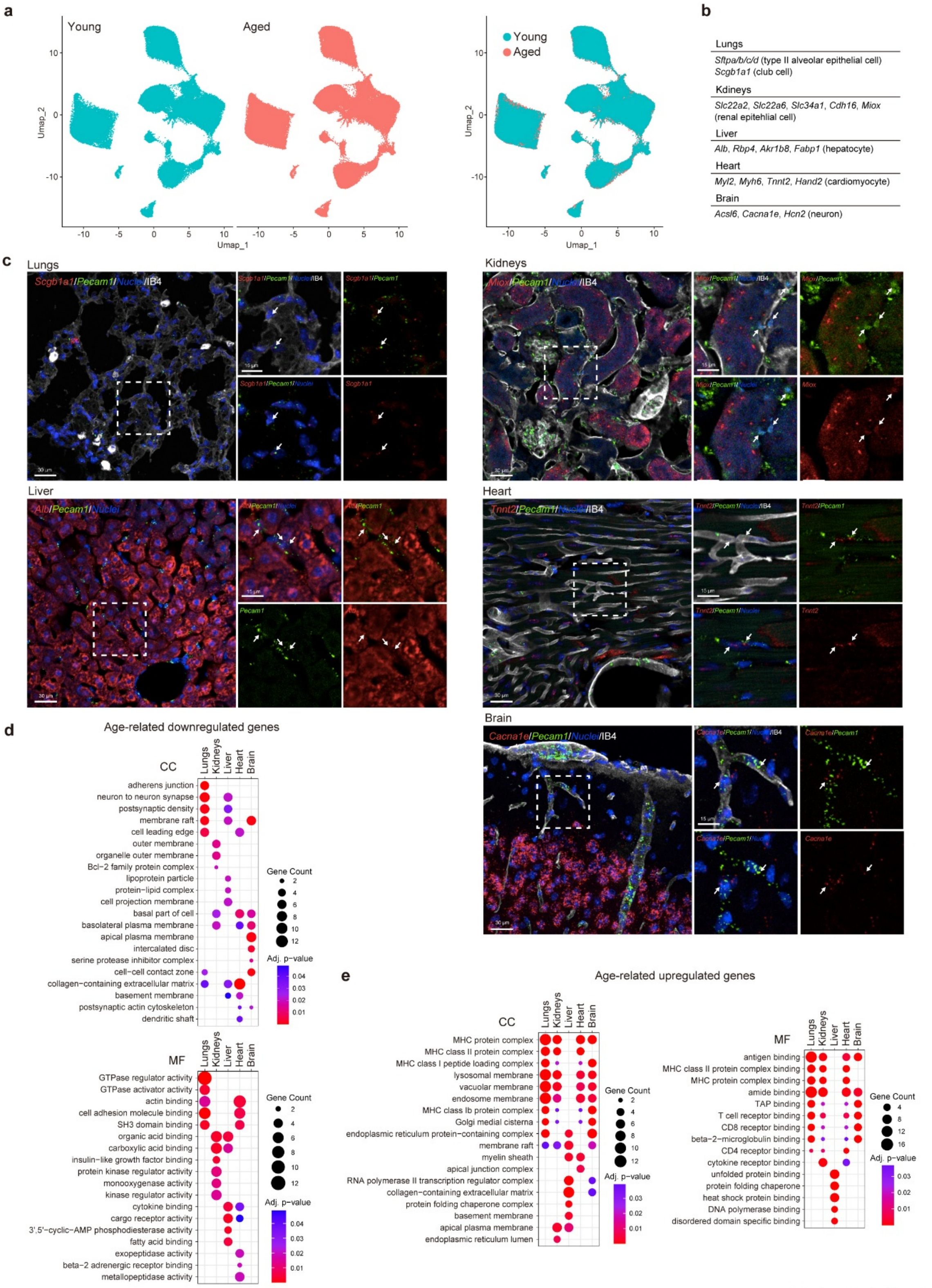

**Extended Fig. 1**

**Extended Fig. 1: Cross-organ endothelial atlas reveals organ-specific gene programs and age-associated transcriptional changes**

**a**, UMAP plots of ECs from young (3-month-old) and aged (24- or 26-month-old) mice,

split by age (left) or merged (right).
**b**, List of genes known to be expressed in parenchymal cells that were also detected in ECs of each organ.
**c**, Representative confocal images of RNAscope in situ hybridization validating the expression of parenchymal cell-associated genes in ECs in all organs harvested from 3-month-old mice. Tissue-specific markers (red): *Scgb1a1* (lungs), *Miox* (kidneys), *Alb* (liver), *Tnnt2* (heart), and *Cacna1e* (brain). *Pecam1* (green) marks ECs. Sections were counterstained with DAPI (nuclei, blue) and IB4 (white). Dashed boxes indicate enlarged regions. Arrows indicate marker-positive ECs.
**d**,**e**, Dot plots showing GO terms in the cellular component (CC) and molecular function (MF) categories enriched in age-downregulated (**d**) and age-upregulated (**e**) genes across organs. Dot size, gene count; dot color, adjusted p-value.
Scale bars, 30 μm (**c**) and 15 μm (enlarged panels in **c**).

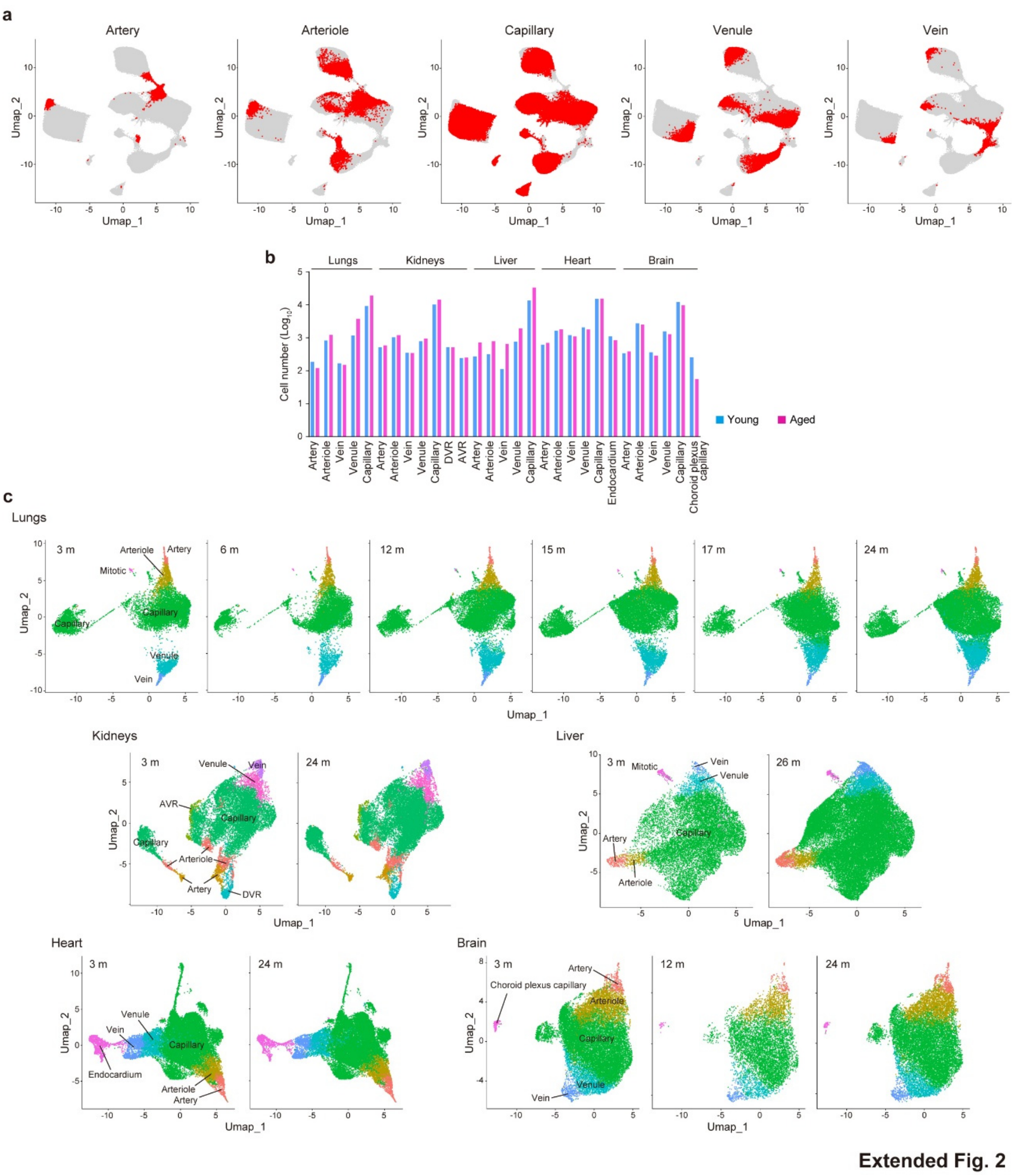

**Extended Fig. 2: Aging induces organ-dependent shifts in vascular subtype composition**

**a**, UMAP plots highlighting the distribution of each vascular subtype (artery, arteriole, capillary, venule, and vein) in all organs examined.

**b**, Bar plots showing cell numbers ($\log_{10}$) of each vascular subtype across organs in young (blue) and aged (pink) mice.

**c**, UMAP plots of ECs with vascular subtype, indicated by color, stratified by age, for each organ. Lungs: 3, 6, 12, 15, 17, and 24 months; kidneys: 3 and 24 months; liver: 3 and 26 months; heart: 3 and 24 months; brain: 3, 12, and 24 months.

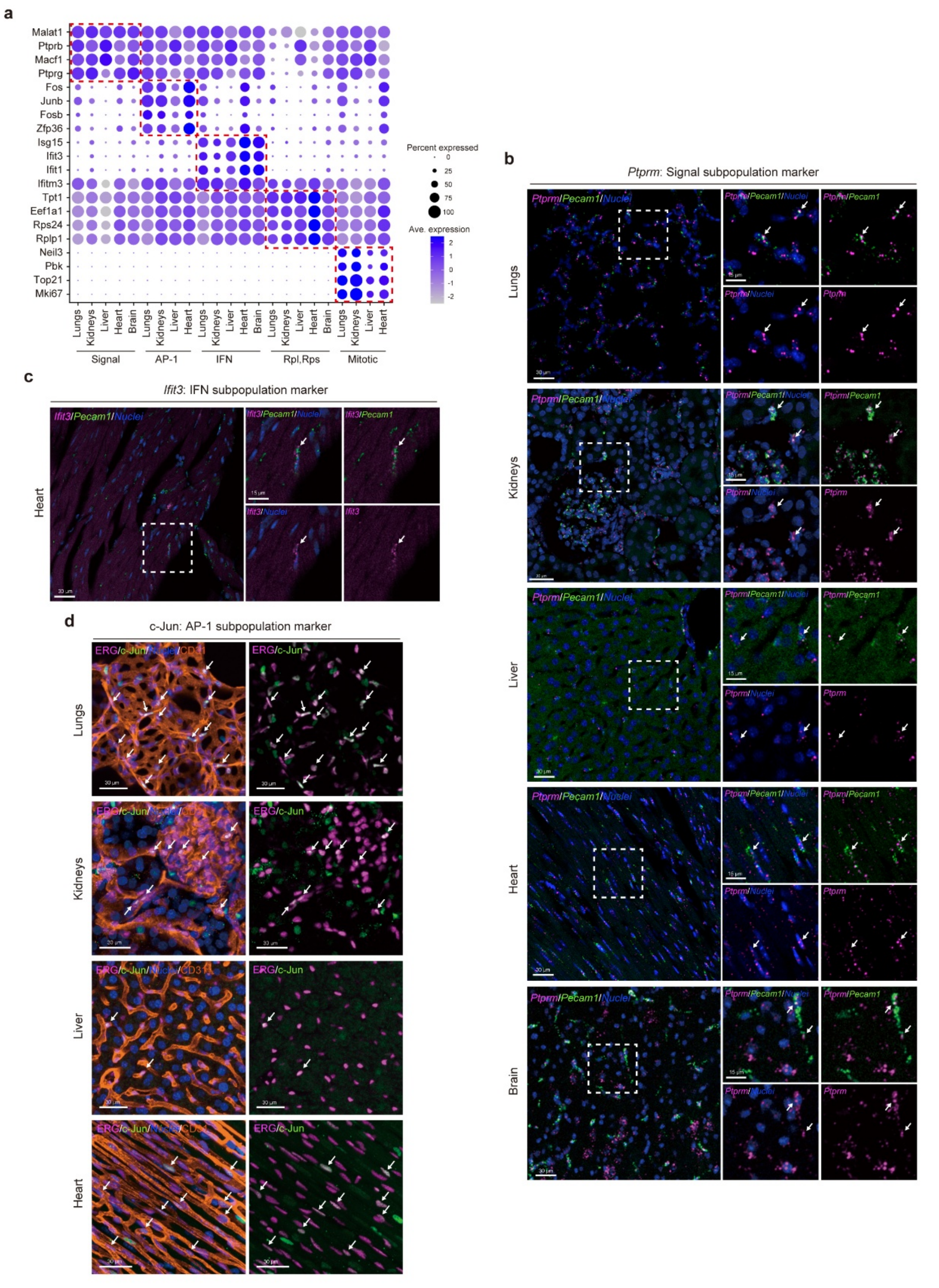

Extended Fig. 3

**Extended Fig. 3: Validation of conserved capillary EC subsets across organs**

**a**, Dot plot showing conserved marker genes of each capillary EC subpopulation across organs. Red dashed boxes highlight subpopulation-specific markers. Dot size, percent expressed; dot color, average expression.

**b**, Representative confocal images of RNAscope in situ hybridization for Ptprm

(magenta, Cap-Signal marker) and Pecam1 (green) with nuclear staining (blue) in the lungs, kidneys, liver, heart, and brain from 12-week-old mice. Dashed boxes indicate regions enlarged in the right panels. Arrows indicate Ptprm-expressing Pecam1$^+$ ECs.
**c**, Representative confocal images of RNAscope in situ hybridization for Ifit3 (magenta, Cap-IFN marker) and Pecam1 (green) with nuclear staining (blue) in the heart brain from 12-week-old mice. Dashed boxes indicate regions enlarged in the right panels. Arrows indicate Ptprm-expressing Pecam1$^+$ ECs.
**d**, Representative confocal images of lungs, liver, heart, and kidneys from 12-week-old mice immunostained for c-Jun (green, Cap-AP-1 marker), ERG (magenta), and CD31 (orange), counterstained with DAPI (nuclei, blue). Arrows indicate c-Jun- and ERG-double-positive ECs.
Scale bars, 30 μm (**b**-**d**) and 15 μm (enlarged panels in **b**, **c**).

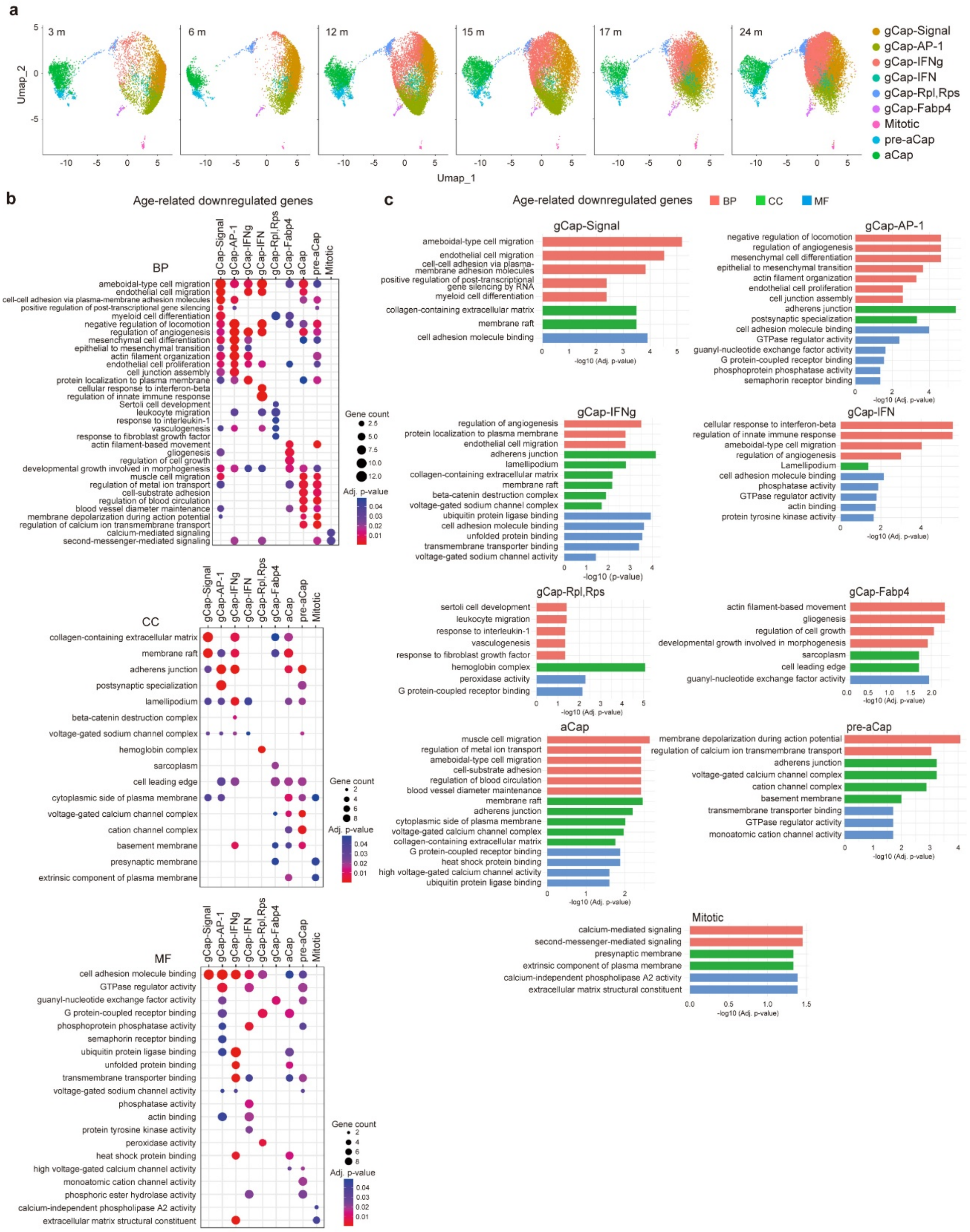

**Extended Fig. 4: Aging progressively erodes homeostatic programs in alveolar capillary EC subpopulations**

**a**, UMAP plots of alveolar capillary ECs, with subpopulations indicated by color, at 3, 6, 12, 15, 17, and 24 months of age.

**b**, Dot plots showing GO terms in the biological process (BP), cellular component (CC), and molecular function (MF) categories enriched in the top 100 age-downregulated

genes across alveolar capillary EC subpopulations. Dot size, gene count; dot color, adjusted p-value.
**c**, Bar plots showing GO terms enriched in the top 100 age-downregulated genes in each alveolar capillary EC subpopulation. x-axis, −log10(adjusted p-value); BP, pink; CC, green; MF, blue.

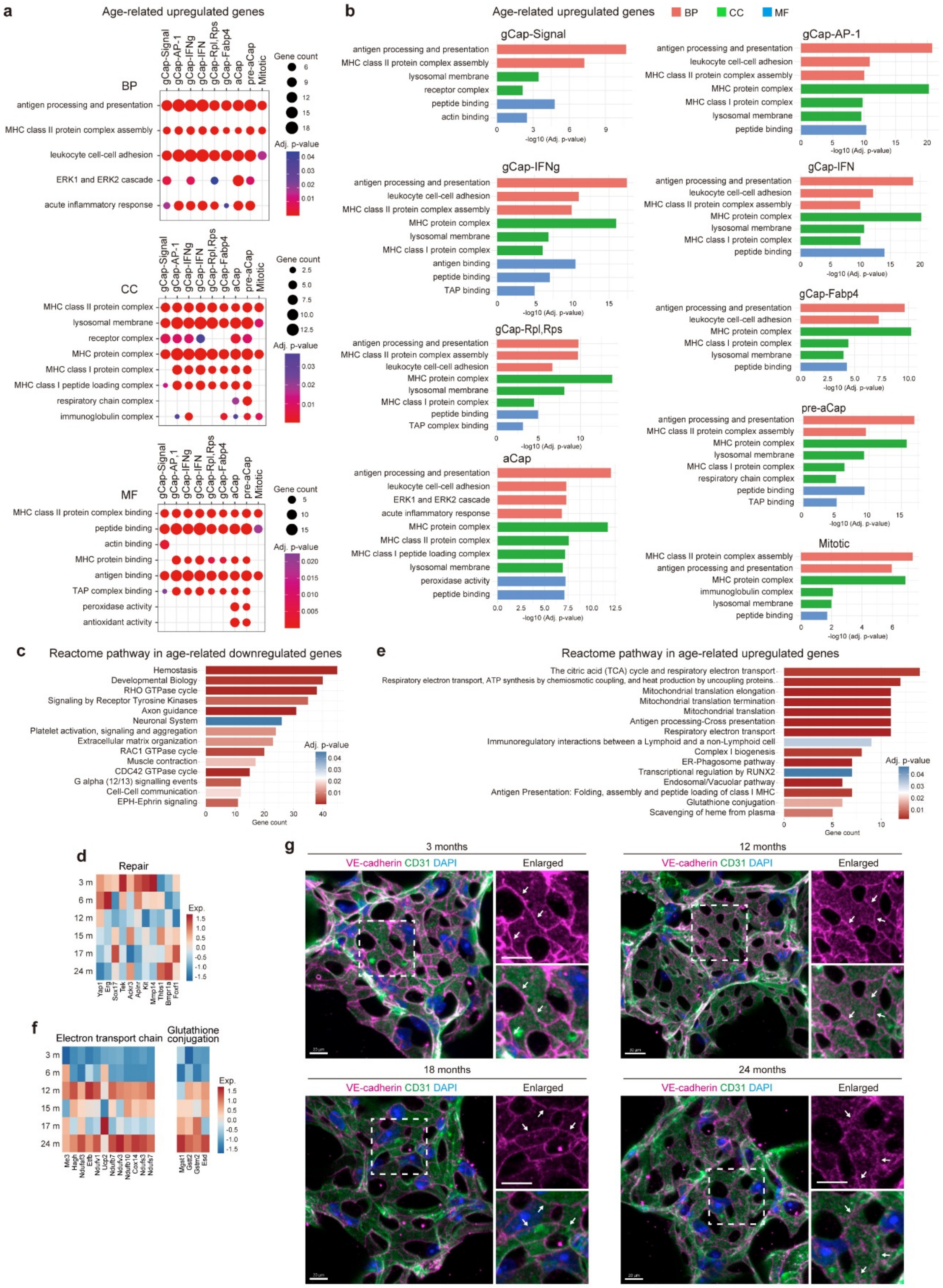

**Extended Fig. 5**

**Extended Fig. 5: Aging induces antigen presentation programs and impairs vascular barrier integrity in alveolar capillary ECs**

**a**, Dot plots showing GO terms in the biological process (BP), cellular component (CC), and molecular function (MF) categories enriched in the top 100 age-upregulated genes across alveolar capillary EC subpopulations. Dot size, gene count; dot color, adjusted p-value.

**b**, Bar plots showing GO terms enriched in the top 100 age-upregulated genes in each alveolar capillary EC subpopulation. x-axis, −log10(adjusted p-value); BP, pink; CC, green; MF, blue.

**c**,**e**, Reactome pathway analysis of the top 100 age-downregulated (**c**) and age-upregulated (**e**) genes in tgCap ECs (gCap-Signal, gCap-AP-1, and gCap-IFNg). x-axis, gene count; color, adjusted p-value.

**d**,**f**, Heatmaps showing normalized expressions (based on z-scores) of repair-related genes (**d**) and electron transport chain- and glutathione conjugation-related genes (**f**) in tgCap ECs at 3, 6, 12, 15, 17, and 24 months of age.

**g**, Representative confocal images of alveoli from lungs of 3-, 12-, 18-, and 24-month-old mice immunostained for VE-cadherin (magenta) and CD31 (green), counterstained with DAPI (blue). Dashed boxes indicate enlarged regions. Arrows indicate VE-cadherin at cell–cell junctions.

Scale bars, 30 μm (**g**) and 10 μm (enlarged panels in **g**).

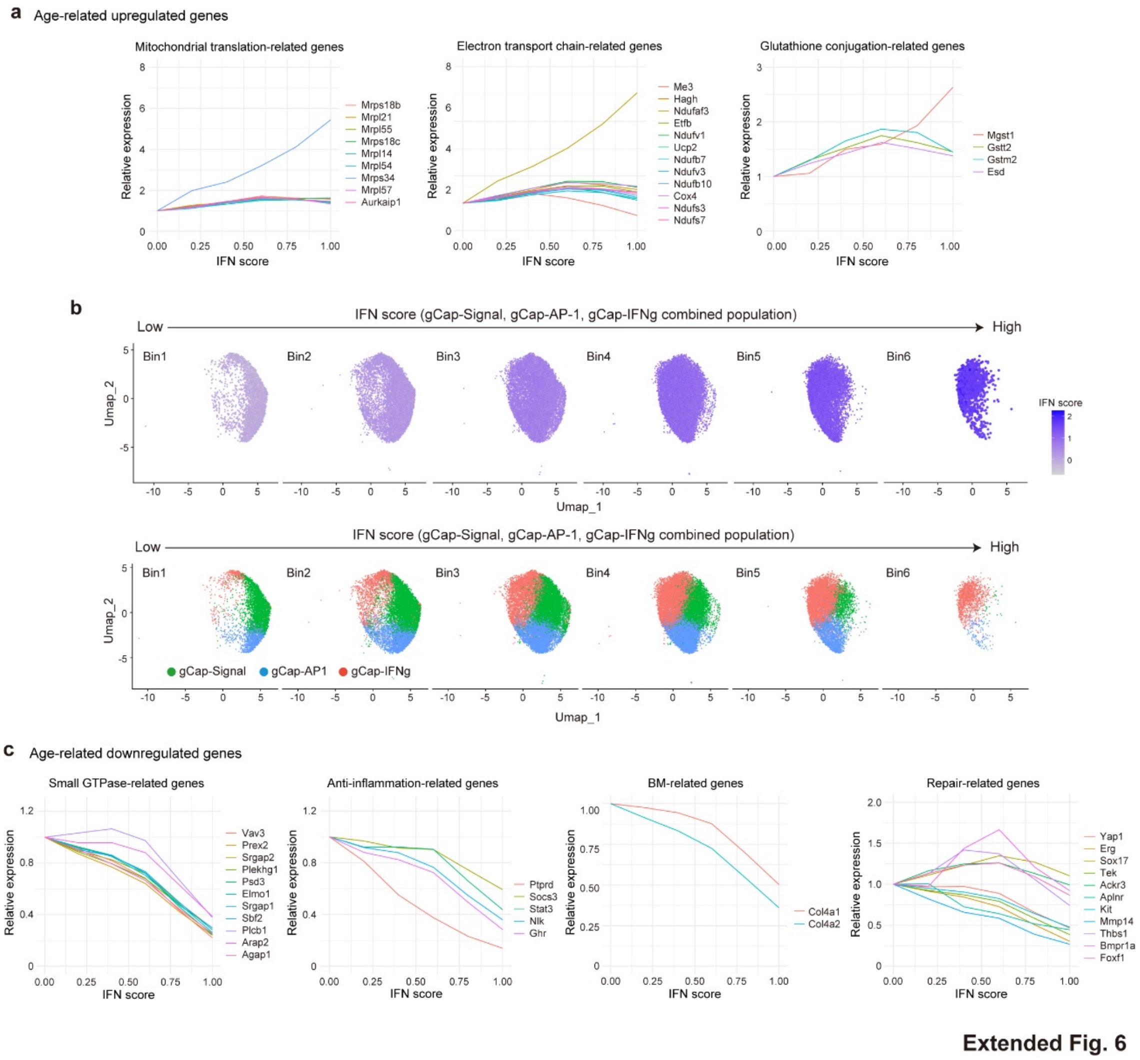

**Extended Fig. 6: The IFN-based aging index captures progressive functional deterioration in alveolar capillary ECs**

**a**, Relative expressions of mitochondrial translation-, electron transport chain-, and glutathione conjugation-related genes in tgCap ECs plotted against the IFN score. Values are normalized to the expression at IFN score = 0.

**b**, UMAP plots of tgCap ECs (gCap-Signal, gCap-AP-1, and gCap-IFNg combined), with IFN scores in color, divided into six bins (upper) or by subpopulation (lower).

**c**, Relative expressions of small GTPase-, anti-inflammatory-, basement membrane-, and repair-related genes in tgCap ECs plotted against the IFN score. Values are normalized to the expression at IFN score = 0.

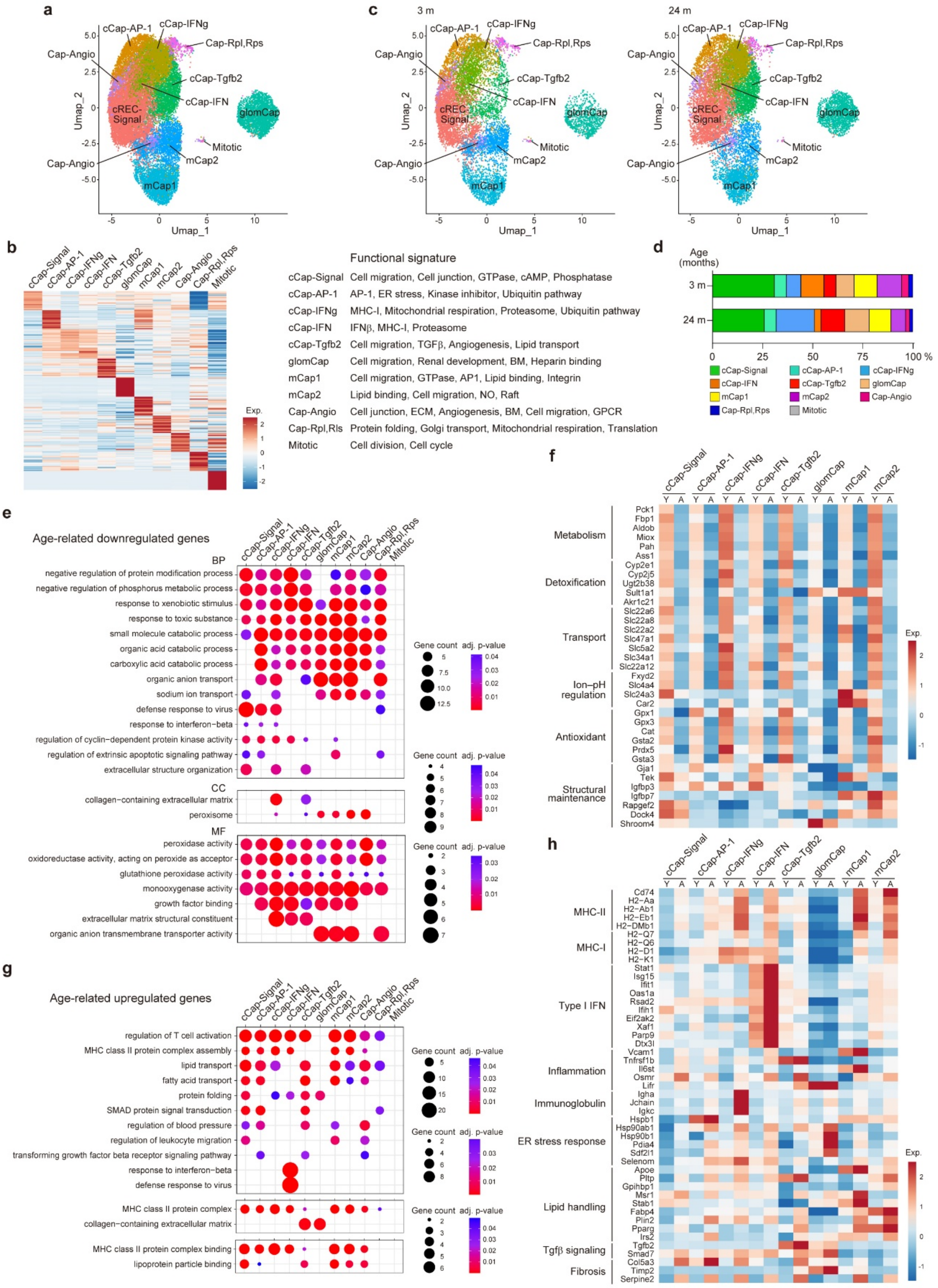

**Extended Fig. 7**

**Extended Fig. 7: Aging remodels renal capillary ECs toward immune-activated and stress-associated states**

**a**, UMAP plot of renal capillary ECs, with subpopulations indicated by color.

**b**, Heatmap showing the top 10 marker genes for each renal capillary EC subpopulation, with functional signatures indicated on the right. Color scale, z-score.
**c**, UMAP plots of renal capillary ECs, with subpopulations indicated by color, stratified by age (3 and 24 months).
**d**, Proportional changes in renal capillary EC subpopulations in young (3-month-old) and aged (24-month-old) mice.
**e**,**g**, Dot plots showing GO terms in the biological process (BP), cellular component (CC), and molecular function (MF) categories enriched in the top 100 age-downregulated (**e**) and age-upregulated (**g**) genes across renal capillary EC subpopulations. Dot size, gene count; dot color, adjusted p-value.
**f**,**h**, Heatmaps showing normalized expressions (based on z-scores) of functionally significant age-downregulated (**f**) and age-upregulated (**h**) genes across renal capillary EC subpopulations in young (Y) and aged (A) mice.

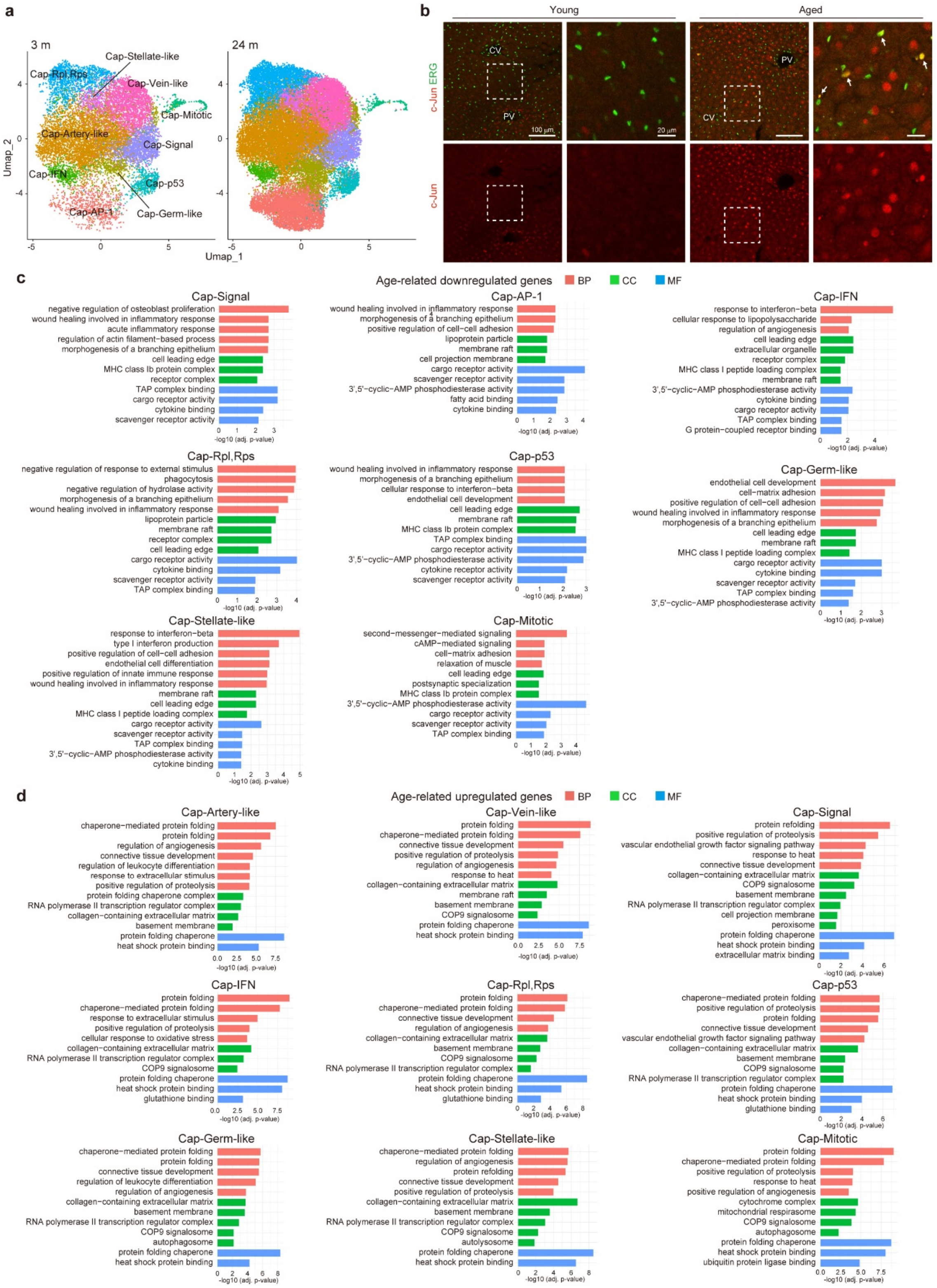

**Extended Fig. 8**

**Extended Fig. 8: Aging expands stress-associated LSECs and attenuates homeostatic programs**

**a**, UMAP plots of hepatic capillary ECs, with subpopulations indicated by color,

stratified by age (3 and 24 months).
**b**, Representative confocal images of liver sections from young and aged mice immunostained for c-Jun (red) and ERG (green). Dashed boxes indicate enlarged regions. Arrows indicate c-Jun- and ERG-double-positive ECs. PV, portal vein; CV, central vein.
**c**,**d**, Bar plots showing GO terms enriched in the top 100 age-downregulated (**c**) and age-upregulated (**d**) genes in each LSEC subpopulation. x-axis, −log10(adjusted p-value); BP, pink; CC, green; MF, blue.
Scale bars, 100 μm (**b**) and 20 μm (enlarged panels in B).

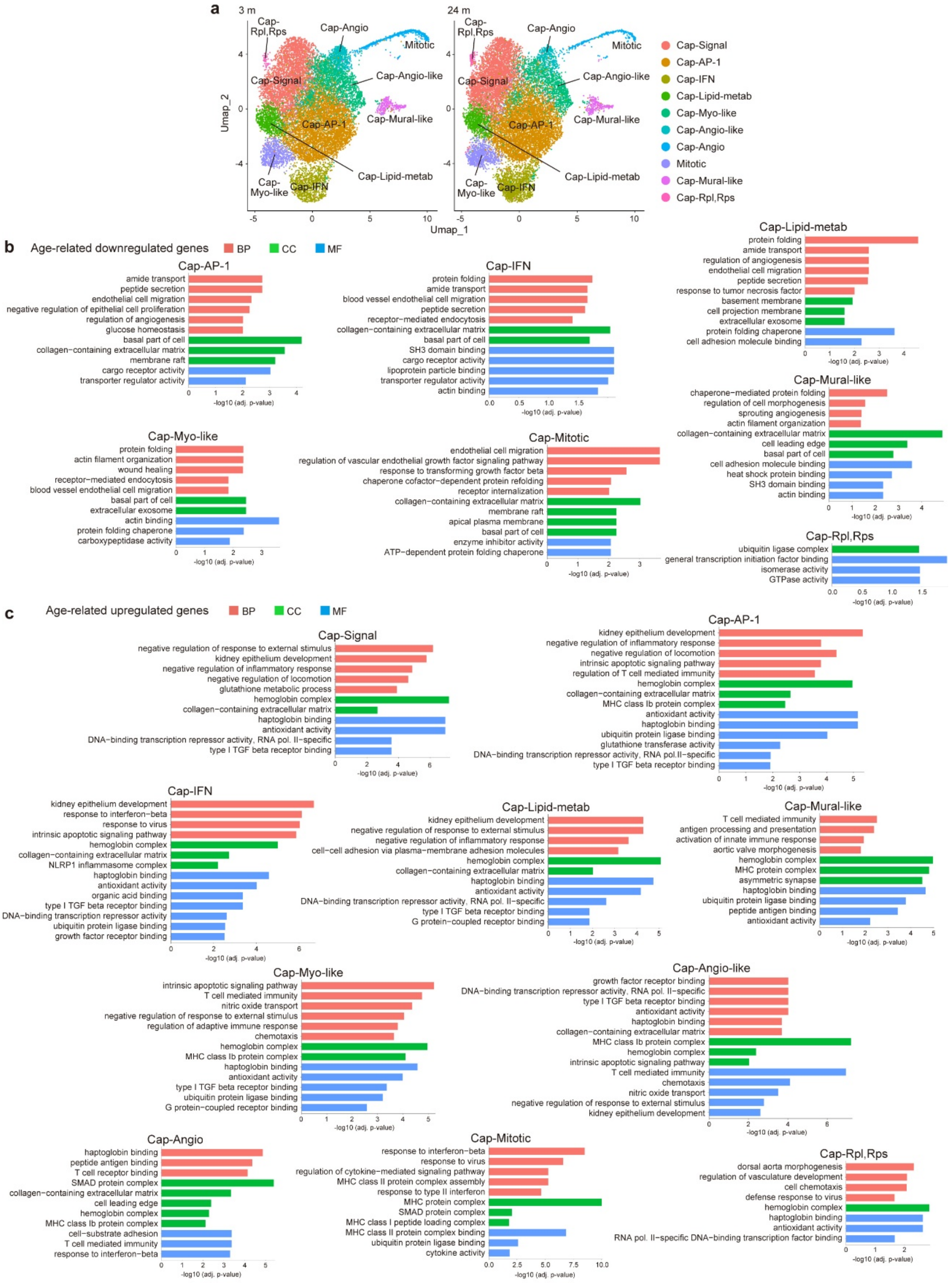

**Extended Fig. 9**

**Extended Fig. 9: Aging impairs homeostatic and angiogenic programs in cardiac capillary ECs**

**a**, UMAP plots of cardiac capillary ECs, with subpopulations indicated by color, stratified by age (3 and 24 months).

**b**,**c**, Bar plots showing GO terms enriched in the top 100 age-downregulated (**b**) and age-upregulated (**c**) genes in each cardiac capillary EC subpopulation. x-axis, −log10(adjusted p-value); BP, pink; CC, green; MF, blue.

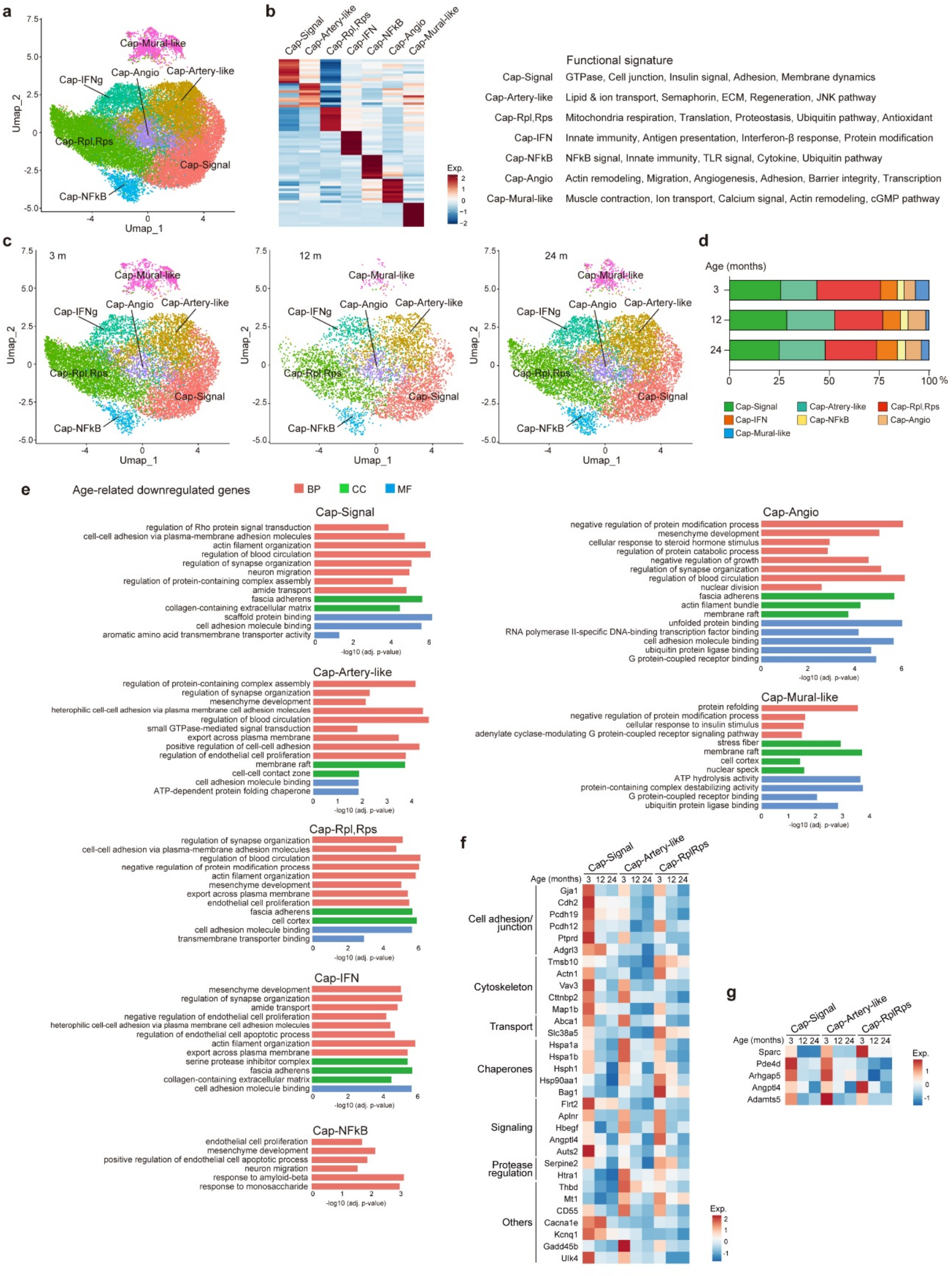

**Extended Fig. 10**

**Extended Fig. 10: Aging attenuates BBB integrity and homeostatic programs in brain capillary ECs**

**a**, UMAP plot of brain capillary ECs, with subpopulations indicated by color.

**b**, Heatmap showing the top 10 marker genes for each brain capillary EC subpopulation,

with functional signatures indicated on the right. Color scale, z-score.
**c**, UMAP plots of brain capillary ECs, with subpopulations indicated by color, stratified by age (3, 12, and 24 months).
**d**, Proportional changes in brain capillary EC subpopulations at 3, 12, and 24 months of age.
**e**, Bar plots showing GO terms enriched in the top 100 age-downregulated genes in each brain capillary EC subpopulation. x-axis, −log10(adjusted p-value); BP, pink; CC, green; MF, blue.
**f**,**g**, Heatmaps showing normalized expressions (based on z-scores) of age-downregulated genes related to the indicated functions (**f**) and BBB dysfunction (**g**) across brain capillary EC subpopulations (Cap-Signal, Cap-Artery-like, and Cap-Rpl,Rps) at 3, 12, and 24 months of age.

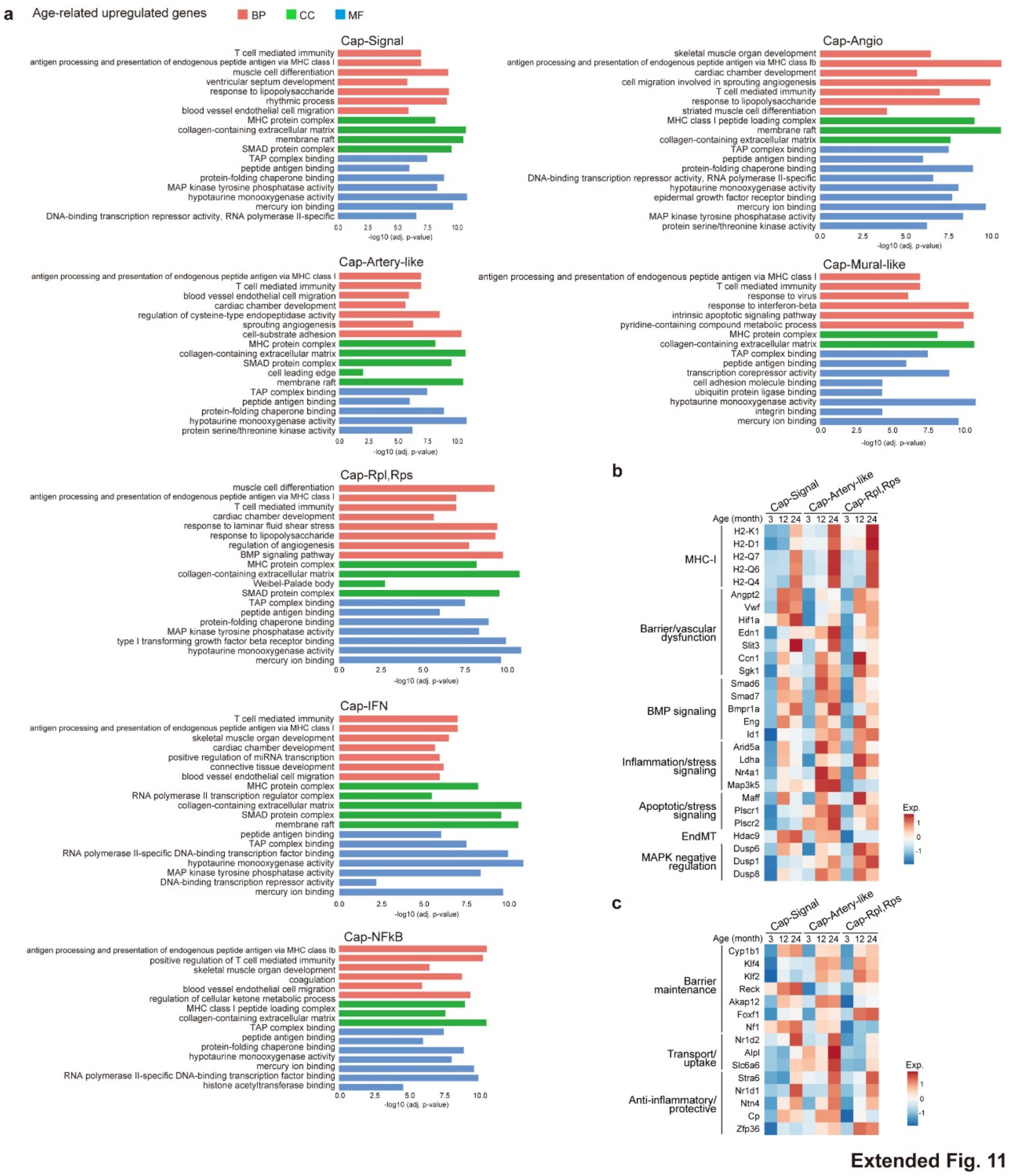

**Extended Fig 11: Aging induces maladaptive and compensatory transcriptional responses in brain capillary ECs**

**a**, Bar plots showing GO terms enriched in the top 100 age-upregulated genes in each brain capillary EC subpopulation. x-axis, −log10(adjusted p-value); BP, pink; CC, green; MF, blue.

**b**,**c**, Heatmaps showing normalized expressions (based on z-scores) of age-upregulated genes related to the indicated functions (**b**) and BBB maintenance and protective functions (**c**) across brain capillary EC subpopulations (Cap-Signal, Cap-Artery-like, and Cap-Rpl,Rps) at 3, 12, and 24 months of age.

#### Supplementary Tables

#### Age-associated erosion of organ-specific endothelial programs compromises tissue function and resilience

Haruko Watanabe-Takano, Tomohiro Ishii, Tomohisa Hayakawa, Hitoshi Iuchi, Hitomi Matsuno, Eri Oguri-Nakamura, Kunihito Arai, Kei Yura, Michiaki Hamada, Daisuke Hishikawa, Shota Toyoshima, Mashito Sakai, Shimpei Higo, Masahiro Morishita, Hirotaka Ishii, Toru Tanaka, Sayo Horibe, Yoshiyuki Rikitake, Taichi Noda, Kimi Araki, Takashi Minami, Shinya Tanaka, Shigetomo Fukuhara

**Table S1. Summary of sequencing and quality control metrics across tissues and samples.**

| Tissue | Sample name | Estimated Number of cells | Mean Reads per cell | Median Genes per Cell | Reads Mapped to Genome(%) | Cell Number after filtering |
|---|---|---|---|---|---|---|
| Lungs | 12w_1 | 7939 | 51,938 | 2,850 | 98.6 | 5928 |
| | 12w_2 | 12,249 | 33,343 | 2,738 | 98.4 | 9589 |
| | 29w_1 | 14,173 | 20,939 | 2,712 | 94.8 | 13269 |
| | 54w_1 | 12,659 | 26,166 | 2,643 | 95.6 | 11581 |
| | 54w_2 | 13,412 | 25,100 | 2,475 | 95.1 | 12120 |
| | 65w_1 | 8,722 | 39,012 | 2,618 | 97.8 | 7867 |
| | 65w_2 | 13,188 | 24,825 | 2,076 | 98 | 11617 |
| | 72w_1 | 11,454 | 36,626 | 2,794 | 98.6 | 8121 |
| | 72w_2 | 13,641 | 29,846 | 2,512 | 98.5 | 11143 |
| | 106w_1 | 5,366 | 61,906 | 2,908 | 97.7 | 4415 |
| | 106w_2 | 18,666 | 18,197 | 1,733 | 97.8 | 17011 |
| | 106w_3 | 11,380 | 31,634 | 2,965 | 97.2 | 9931 |
| Kidneys | 11w_1 | 15,308 | 28,543 | 1,952 | 96.9 | 10996 |
| | 11w_2 | 11,818 | 35,985 | 1,882 | 97 | 8797 |
| | 106w_1 | 14,024 | 31,779 | 1,759 | 96.8 | 10402 |
| | 106w_2 | 13,763 | 30,438 | 1,929 | 96.9 | 9593 |
| Liver | LiverEC_3mo_1 | 15,661 | 27,821 | 3,019 | 94.9 | 15,018 |
| | LiverEC_26mo_1 | 20,650 | 21,263 | 2,823 | 95.9 | 19,851 |
| | LiverEC_26mo_2 | 18,372 | 23,990 | 3,023 | 96.4 | 17,662 |
| Heart | 190-1_3month-1 | 9,562 | 48,880 | 2,276 | 98.6 | 9,065 |
| | 190-2_24month-1 | 6,652 | 70,158 | 2,200 | 98.7 | 6,357 |
| | 190-3_3month-2 | 8,170 | 42,219 | 2,309 | 98.8 | 7,694 |
| | 190-4_24month-2 | 8,165 | 47,911 | 2,316 | 98.6 | 7,972 |
| | 190-9_3month-3 | 6,628 | 59,786 | 2,637 | 97.8 | 6,404 |
| | 190-10_24month-3 | 8,893 | 40,721 | 2,466 | 97.7 | 8,611 |
| Brain | 191-1_3month-1 | 11,860 | 35,765 | 2,444 | 98.1 | 7,755 |
| | 191-4_24month-2 | 6,925 | 60,563 | 3,016 | 97.7 | 6,108 |
| | 191-5_12month-1 | 7,178 | 62,642 | 2,951 | 98.5 | 7,016 |
| | 191-6_24month-3 | 8,997 | 50,178 | 2,894 | 98.4 | 8,820 |
| | 191-7_3month-3 | 7,893 | 53,657 | 2,997 | 98.3 | 7,755 |
| Median | | 11636 | 35,875 | 2,628 | 97.8 | 8942.5 |

**Table S2. Dataset-specific filtering thresholds used for quality control.**

| Tissue | Sample name | min.cells | min.features | nFeatures | percent_mt |
|---|---|---|---|---|---|
| Lungs | 12w_1 | 3 | 200 | 200<nFeatures<6000 | 0<percent_mt<7 |
| | 12w_2 | 3 | 200 | 200<nFeatures<6000 | 0<percent_mt<7 |
| | 29w_1 | 3 | 200 | 200<nFeatures<6000 | 0<percent_mt<7 |
| | 54w_1 | 3 | 200 | 200<nFeatures<6000 | 0<percent_mt<7 |
| | 54w_2 | 3 | 200 | 200<nFeatures<6000 | 0<percent_mt<7 |
| | 65w_1 | 3 | 200 | 200<nFeatures<6000 | 0<percent_mt<7 |
| | 65w_2 | 3 | 200 | 200<nFeatures<6000 | 0<percent_mt<7 |
| | 72w_1 | 3 | 200 | 200<nFeatures<6000 | 0<percent_mt<7 |
| | 72w_2 | 3 | 200 | 200<nFeatures<6000 | 0<percent_mt<7 |
| | 106w_1 | 3 | 200 | 200<nFeatures<6000 | 0<percent_mt<7 |
| | 106w_2 | 3 | 200 | 200<nFeatures<6000 | 0<percent_mt<7 |
| | 106w_3 | 3 | 200 | 200<nFeatures<6000 | 0<percent_mt<10 |
| Kidneys | 11w_1 | 3 | 200 | 200<nFeatures<6000 | 0<percent_mt<10 |
| | 11w_2 | 3 | 200 | 200<nFeatures<6000 | 0<percent_mt<10 |
| | 106w_1 | 3 | 200 | 200<nFeatures<6000 | 0<percent_mt<10 |
| | 106w_2 | 3 | 200 | 200<nFeatures<6000 | 0<percent_mt<10 |
| Liver | LiverEC_3mo_1 | | | 200<nFeatures<6000 | 0<percent_mt<8 |
| | LiverEC_26mo_1 | | | 200<nFeatures<6000 | 0<percent_mt<8 |
| | LiverEC_26mo_2 | | | 200<nFeatures<6000 | 0<percent_mt<8 |
| Heart | 190-1_3month-1 | | | 500<nFeatures<6000 | percent_mt<10 |
| | 190-2_24month-1 | | | 500<nFeatures<6000 | percent_mt<10 |
| | 190-3_3month-2 | | | 500<nFeatures<6000 | percent_mt<10 |
| | 190-4_24month-2 | | | 500<nFeatures<6000 | percent_mt<10 |
| | 190-9_3month-3 | | | 500<nFeatures<6000 | percent_mt<10 |
| | 190-10_24month-3 | | | 500<nFeatures<6000 | percent_mt<10 |
| Brain | 191-1_3month-1 | | | 500<nFeatures<6000 | percent_mt<10 |
| | 191-4_24month-2 | | | 500<nFeatures<6000 | percent_mt<10 |
| | 191-5_12month-1 | | | 500<nFeatures<6000 | percent_mt<10 |
| | 191-6_24month-3 | | | 500<nFeatures<6000 | percent_mt<10 |
| | 191-7_3month-3 | | | 500<nFeatures<6000 | percent_mt<10 |

**Table S3. Marker genes used for cluster annotation across tissues.**

| Lungs | Marker genes | | | | |
|---|---|---|---|---|---|
| Endothelial cells | Pecam1 | Emcn | Cdh5 | Icam2 | Cldn5 |
| Pericytes | Pdgfrb | | | | |
| Lymphatic endothelial cells | Prox1 | | | | |
| Hematopoetic cells | Ptprc | | | | |
| Fibroblasts | Pdgfra | | | | |
| Smooth muscle cells | Acta2 | | | | |
| Type I alveolar epithelial cells | Ager | | | | |
| Type II alveolar epithelial cells | Sftpc | Sftpd | Nkx2-1 | | |
| Epithelial cells | Epcam | Scgb1a1 | | | |

| Kidneys | Marker genes | | | | |
|---|---|---|---|---|---|
| Endothelial cells | Pecam1 | Emcn | Cdh5 | Icam2 | Cldn5 |
| Pericytes | Pdgfrb | | | | |
| Lymphatic endothelial cells | Prox1 | Lyve1 | | | |
| Hematopoetic cells | Ptprc | | | | |
| Fibroblasts | Pdgfra | | | | |
| Tubular epithelial cells | Slc27a2 | Lrp2 | Slc12a1 | Aqp2 | Hsd11b2 |
| Podicytes | Nphs1 | Nphs2 | | | |
| Mesangial cells | Myl9 | Ren1 | | | |

| Liver | Marker genes | | | |
|---|---|---|---|---|
| Endothelial cells | Pecam1 | Cdh5 | | |
| Hapatocyte | Alb | Ttr | Apoa1 | Apoa2 |
| Lymph | Prox1 | | | |
| Kupffer | Adgre1 | C1qa | C1qc | |

| Heart | Marker genes | | | | |
|---|---|---|---|---|---|
| Endothelial cells | Pecam1 | Emcn | Cdh5 | Kdr | Cldn5 |
| Lymphatic | Flt4 | Prox1 | Pdpn | Lyve1 | |
| Pericytes | Pdgfrb | Abcc9 | Kcnj8 | | |
| Smooth muscle cells | Acta2 | Myh11 | | | |
| Fibroblasts | Col1a1 | Fbln | Pdgfra | Tcf21 | Pdgfra |
| Erythrocytes | Hbb-bt | Hba-a1 | Hba-a2 | | |
| Platelet | Itga2b | Pf4 | Fcer1g | | |
| Macrophage | Cd68 | Csf1 | Mpeg1 | | |
| T cell | Cd52 | Cd53 | Coro1a | | |
| Myocyte | Actc1 | Mb | Myl2 | Tnnt2 | |

| Brain | Marker genes | | | | |
|---|---|---|---|---|---|
| Endothelial cells | Cldn5 | Cdh5 | Pecam1 | Pdgfb | Kdr |
| Pericytes | Pdgfrb | Abcc9 | Higd1b | Kcnj8 | |
| Smooth muscle cells | Myh11 | Acta2 | Tagln | | |
| Microglia | Ctss | Cx3cr1 | Aif1 | | |
| Macrophage | Csf1 | Cd68 | Mpeg1 | C1qa | |
| Ependymal cells | Htr2c | Enpp2 | Kcnj13 | Ecrg4 | |

**Table S4. Marker genes used for level 2 endothelial cell sub-clustering and vascular subtype annotation across tissues.**

| Lungs | | | | |
|---|---|---|---|---|
| Artery | Gja5 | Bmx | Mgp | Eln |
| Arteriole | Atp13a3 | Plac8 | Cxcl12 | |
| Capillary | Kit | Glp1r | Scn7a | Car4 |
| Venule | Slc6a2 | Prss23 | | |
| Vein | Vwf | Nr2f2 | Eln | |
| Mitotic | Top2a | MKi67 | | |

| Kidneys | Marker genes | | | | |
|---|---|---|---|---|---|
| Artery | Mgp | Eln | Fbln2 | | |
| Arteriole | Sox17 | S100a6 | Tgfb2(gromellular) | Tspan8(gromellular) | |
| Capillary | Npr3 | Igfbp5 | Kdr | Lpl(gromellular) | |
| Venule | Tnxb | | | | |
| Vein | Bgn | | | | |
| Mitotic | Top2a | MKi67 | | | |
| Ascending Vasa Recta(AVR) | Fxyd6 | Cryab | Fxyd2 | | |
| Descending Vasa Recta(DVR) | Aqp1 | Slc14a1 | S100a4 | Scin | Ifi2712a |

| Liver | Marker genes | | | |
|---|---|---|---|---|
| Artery | Vwf | Sdc1 | Adgrg6 | |
| Arteriole | Msr1 | Adgrg6 | Stab2 | |
| Capillary | Stab2 | Lyve1 | Ctsl | |
| Venule | Kit | Wnt2 | Stab2 | |
| Vein | Vwf | Rspo3 | Wnt2 | Kit |
| Mitotic | Pcna | Top2a | | |

| Heart | Marker genes | | | | |
|---|---|---|---|---|---|
| Artery | Gja4 | Gja5 | Bmx | Jag1 | Sox17 |
| Arteriole | Glul | Hey1 | | | |
| Capillary | Car4 | Aqp7 | Rgcc | | |
| Venule | Aplnr | | | | |
| Vein | Ephb4 | Nr2f2 | | | |
| Endocardium | Col3a1 | Npr3 | Plvap | | |

| Brain | Marker genes | | | |
|---|---|---|---|---|
| Artery | Sema3g | Vegfc | Bmx | Gkn3 |
| Arteriole | Glul | Slc26a10 | Tgfb2 | |
| Capillary | Mfsd2a | Rgcc | | |
| Venule | Tfrc | Slc16a1 | | |
| Vein | Nr2f2 | Slc38a5 | | |
| Choroid plexsus capillary | Plvap | Plpp1 | | |

**Table S5. Conserved marker genes used for level 3 capillary sub-clustering across tissues, corresponding to Figure S3A.**

| Signal | AP-1 | IFN | Rpl,Rps |
|---|---|---|---|
| Malat1 | Fos | Isg15 | Tpt1 |
| Ptprb | Junb | Ifit3 | Eef1a1 |
| Macf1 | Fosb | Ifit1 | Rps24 |
| Ptprg | Zfp36 | Ifitm3 | Rplp1 |
| Psd3 | Atf3 | Rsad2 | Fau |
| Arl15 | Ier3 | Irf7 | Rpl19 |
| Ptk2 | Socs3 | Ifi47 | Rpl27a |
| Arhgap29 | Jun | Ifit3b | Rps14 |
| Itpkb | Gadd45g | Rtp4 | Fth1 |
| Arhgap31 | Btg2 | Xaf1 | Rpl41 |
| Zfp608 | Jund | Cmpk2 | Ptma |
| Rbms1 | Ier5 | Bst2 | Rpl23 |
| Tek | Ier2 | Gbp7 | Rpl28 |
| Tmtc2 | Nfkbiz | Iigp1 | Rps4x |
| Tjp1 | Klf4 | Lgals9 | Rpl9 |
| Pnisr | H3f3b | Ifi35 | Rps27 |
| Sema6a | Dusp1 | Usp18 | Ftl1 |
| Utrn | Eif1 | Ifi44 | Rps16 |
| St6galnac3 | Rhob | Oasl2 | Rps19 |
| Zcchc7 | Btg1 | Ly6a | Rpl30 |
| Ica1 | Ppp1r15a | Ly6e | Rps27a |
| Setd5 | Plk2 | Psmb9 | Rpl17 |
| Ankrd44 | Cebpd | Igtp | Rps8 |
| Myo6 | Ubb | B2m | Rpl26 |
| Atrx | Sqstm1 | Irgm1 | Rps11 |

**Table S6. Ribosomal protein and sex-specific genes excluded prior to GO analysis.**

| Ribosomal_protein_genes | Sex-specific genes |
|---|---|
| Rpl7 | Xist |
| Rpl31 | Ddx3y |
| Rpl37a | |
| Rps6kc1 | |
| Rpl7a | |
| Rpl12 | |
| Rpl35 | |
| Rps21 | |
| Rpl22l1 | |
| Rps3a1 | |
| Rps27 | |
| Rpl34 | |
| Rps20 | |
| Rps6 | |
| Rps8 | |
| Rps6ka1 | |
| Rpl11 | |
| Rpl22 | |
| Rpl9 | |
| Rpl5 | |
| Rplp0 | |
| Rpl6 | |
| Rpl21 | |
| Rpl32 | |
| Rps9 | |
| Rpl28 | |
| Rps5 | |
| Rps19 | |
| Rps16 | |
| Rps11 | |
| Rpl13a | |
| Rpl18 | |
| Rps17 | |
| Rps3 | |
| Rpl27a | |
| Rps13 | |
| Rps15a | |
| Rplp2 | |
| Rpl18a | |
| Rpl13 | |
| Rps25 | |
| Rpl10-ps3 | |
| Rplp1 | |
| Rpl4 | |
| Rps27l | |
| Rpl29 | |
| Rps27rt | |
| Rpsa | |
| Rpl14 | |
| Rps12 | |
| Rps15 | |
| Rpl41 | |
| Rps26 | |
| Rps27a | |
| Rpl26 | |
| Rpl23a | |
| Rpl9-ps1 | |
| Rps6kb1 | |
| Rpl23 | |
| Rpl19 | |
| Rpl27 | |
| Rpl38 | |
| Rps7 | |
| Rpl10l | |
| Rps29 | |
| Rpl36al | |
| Rps6kl1 | |
| Rps6ka5 | |
| Rps23 | |
| Rpl15 | |
| Rps24 | |
| Rpl36a-ps1 | |
| Rpl37 | |
| Rpl30 | |
| Rpl8 | |
| Rpl3 | |
| Rps19bp1 | |
| Rpl39l | |
| Rpl35a | |
| Rpl24 | |
| Rps6ka2 | |
| Rps2 | |
| Rpl3l | |
| Rps10 | |
| Rpl10a | |
| Rps28 | |
| Rps18 | |
| Rpl7l1 | |
| Rpl36 | |
| Rpl36-ps4 | |
| Rps14 | |
| Rpl17 | |
| Rps6kb2 | |
| Rps6ka4 | |
| Rpl9-ps6 | |
| Rpl39 | |
| Rpl10 | |
| Rps4x | |
| Rps6ka6 | |
| Rpl36a | |
| Rps6ka3 | |

**Table S7. Aging index-defining genes used to calculate module scores across tissues and species.**

| Lungs | Liver | Heart | Lungs Human |
|---|---|---|---|
| Cd74 | Icam1 | Smad6 | CD74 |
| B2m | Vcam1 | Smad7 | B2M |
| Tap1 | | Fgf2 | TAP1 |
| Mrps34 | | Angpt2 | MRPS34 |
| Ndufaf3 | | Edn1 | NDUFAF3 |
| H2-K1 | | Robo2 | HLA-DMB |
| H2-D1 | | Cxcl12 | HLA-DRB1 |
| H2-Aa | | | |
| H2-DMb1 | | | |
| H2-Eb1 | | | |
